# Identification of translation events that drive nonsense-mediated mRNA decay reveals functional roles for noncoding RNAs

**DOI:** 10.1101/2025.08.15.670413

**Authors:** David J. Young, Yuejun Wang, Nicholas R. Guydosh

## Abstract

The nonsense-mediated mRNA decay (NMD) pathway targets mRNAs undergoing premature translation termination for degradation. Previously, RNA-seq of yeast lacking NMD revealed that most genes targeted by NMD lack obvious premature termination codons (PTCs). We developed a combined approach using RNA-seq and a novel 40S ribosome profiling strategy to identify cryptic premature termination events that could account for NMD on nearly all these transcripts, including many non-coding RNA transcript isoforms associated with annotated genes. Many NMD-targeted transcripts appear to be involved in two-promoter gene regulatory systems and share properties with long un-decoded transcript isoforms (LUTIs). In particular, we show that the *DAL5* LUTI regulates expression of the *DAL5* protein-coding mRNA in response to changes to environmental nitrogen. Our work expands the functional roles for LUTIs and establishes the importance of NMD in their regulation.

**HIGHLIGHTS:**

1. 40S ribosome profiling reveals a comprehensive catalog of NMD substrates in yeast.
2. Most NMD targets have short uORFs or iORFs, resulting in premature termination.
3. Many long undecoded transcript isoforms (LUTIs) contain NMD-triggering uORFs.
4. LUTIs, such as *DAL5*, play a role in metabolic regulation (i.e. nitrogen and thiamine).

## INTRODUCTION

The nonsense-mediated mRNA decay (NMD) pathway degrades mRNAs undergoing premature translation termination in eukaryotes (He and Jacobson, 2015; Losson and Lacroute, 1979; Maquat et al., 1981). Genes encoding the core components of the NMD pathway were initially identified in *Saccharomyces cerevisiae* – *NAM7* (*UPF1*), *NMD2* (*UPF2*), and *UPF3* (Cui et al., 1995; He and Jacobson, 1995; Lee and Culbertson, 1995; Leeds et al., 1991; Leeds et al., 1992). Despite extensive research, the exact biological function of NMD remains poorly understood. A compelling model for recognition of PTCs is the “faux-UTR model” that links the efficiency of termination to NMD (Amrani et al., 2004; He and Jacobson, 2015). Efficient translation termination depends on interactions between the translation termination release factors (eRF1 and eRF3) and proteins bound to the 3’-UTR and polyA tail. In the case of a premature termination (stop) codon (PTC), the ribosome is positioned too far away from the 3’-UTR and polyA tail for these interactions to occur, which allows the NMD machinery to associate with the ribosome and initiate NMD (Amrani *et al*., 2004; Andjus et al., 2024; Bharti et al., 2024; Decourty et al., 2014; Kebaara and Atkin, 2009; Kolakada et al., 2025; Mangkalaphiban et al., 2021; Meydan and Guydosh, 2020; Muhlrad and Parker, 1999; Wu et al., 2020). In some eukaryotes, detection of PTCs can be enhanced by downstream exon-junction complexes (Hogg and Goff, 2010; Le Hir et al., 2001; Singh et al., 2008) or inhibited by other proteins (Fritz et al., 2020; Kishor et al., 2019). Nevertheless, the core NMD pathway is conserved in *S. cerevisiae*, making it a tractable organism for addressing basic questions of NMD function and mechanism (Munoz et al., 2023).

Several structural classes of NMD targets have been identified in yeast based on their stabilization upon inactivation of the NMD pathway, including: (1) genes with mutations that introduce nonsense codons, including pseudogenes (He et al., 2003; Losson and Lacroute, 1979; Peltz et al., 1993), (2) mRNAs that have retained their introns either due to splicing errors or developmental programming e.g., *RPL28* (*CYH2*) and *REC107* (*MER2*) (He et al., 1993), (3) mRNAs containing upstream open reading frames (uORFs) in their 5’UTR e.g., *CPA1* (Arribere and Gilbert, 2013; Gaba et al., 2005; He *et al*., 2003; May et al., 2023; Oliveira and McCarthy, 1995; Ruiz-Echevarria and Peltz, 2000), (4) mRNAs with long 3’ UTRs e.g. *PGA1* and *COX19* (Kebaara and Atkin, 2009; Muhlrad and Parker, 1999; Peccarelli et al., 2016), and (5) mRNAs that utilize programmed +1 frameshifting in their translation e.g., *EST3* (He *et al*., 2003).

Only some mRNAs with uORFs are targeted by the NMD pathway (Gaba *et al*., 2005; He *et al*., 2003; Ruiz-Echevarria and Peltz, 2000). At least some that are resistant to NMD appear to exhibit efficient termination (Gaba *et al*., 2005), potentially due to sequences in proximity to the uORF stop codon that recruit factors that facilitate termination e.g., Pub1 (Ruiz-Echevarria and Peltz, 2000). Similarly, only some mRNAs with long 3’UTRs are targeted to NMD. The basis of this resistance is unclear though it may depend on the length of the main ORF (Decourty *et al*., 2014). In addition, the level of ribosome queuing upstream of a stop codon or the identity of the codon immediately prior to the stop codon (“penultimate codon”) have been proposed to affect termination efficiency and could therefore be important (Bharti *et al*., 2024; Kolakada *et al*., 2025; Mangkalaphiban *et al*., 2021; Meydan and Guydosh, 2020). Studies have also shown that the level of readthrough at stop codons can affect NMD (Pacheco et al., 2024). Apparent NMD sensitivity is also affected by the efficiency of competing mRNA turnover pathways (Mabin et al., 2025). Given the above, it remains a challenge to predict how effective any particular PTC is at promoting degradation.

Aside from the annotated genome, many unannotated transcripts have been identified as NMD substrates in yeast. These include stable unannotated transcripts (SUTs; (Xu et al., 2009) and RNAs stabilized in the absence of genes encoding specific RNA decay factors, including the cytoplasmic 5’-3’ RNA exonuclease *XRN1* (XUTs; (van Dijk et al., 2011; Wery et al., 2016) and the catalytic subunit of the decapping complex *DCP2* (Geisler et al., 2012). It has been shown that 70% of XUT transcripts are NMD substrates (Malabat et al., 2015; Wery *et al*., 2016) and another study reported that of ∼800 unannotated transcripts identified, 16% of were NMD-sensitive (Smith et al., 2014).

A recent exhaustive effort defined NMD targets in yeast by performing RNA-seq on WT, *upf1*Δ, *upf2*Δ, and *upf3*Δ yeast strains (Celik et al., 2017). Among protein-coding genes and functional RNAs, 88% of NMD-targeted transcripts appeared to lack an obvious PTC that would fit into one of the five structural classes described above. These are likely to be direct targets of NMD because prior studies showed that RNAs stabilized in *upf*Δ strains directly associate with Upf1 protein (Johansson et al., 2007). The genes without a recognizable PTC were generally translated inefficiently, suggesting that alternative translation events could be responsible for NMD (Celik *et al*., 2017). Alternative NMD-targeting translation events have been reported, and many relate to an often-underappreciated observation that multiple transcript isoforms can be associated with a given yeast gene (Miura et al., 2006; Zhang and Dietrich, 2005). Whereas older studies using RNA-seq and tiling microarrays limited each gene to only one TSS (Nagalakshmi et al., 2008; Xu *et al*., 2009), more recent studies using TIF-Seq, which sequences the 5’ and 3’ ends of each transcript, and TL-seq, a high-throughput version of 5’ Rapid Amplification of cDNA Ends (RACE), confirmed multiple TSSs for a majority of *S. cerevisiae* genes (Arribere and Gilbert, 2013; Malabat *et al*., 2015; Pelechano et al., 2013). These later studies identified several additional classes of transcript isoforms that are targeted to NMD, including one where two or more distinct TSSs, which were at least 50 nucleotides apart, exist for at least 6.3% of genes (Arribere and Gilbert, 2013).

Large numbers of two-transcript gene regulatory systems where transcription generates a 5’-extended RNA represses the promoter of a downstream canonical transcript have recently been described (Chen et al., 2017; Cheng et al., 2018; Chia et al., 2017; Tresenrider et al., 2021; Van Dalfsen et al., 2018). Transcription generates these 5’-extended transcripts, known as long <u>u</u>ndecoded transcript isoforms (LUTIs), results in repressive chromatin marks being deposited on the downstream (proximal) promoter (Chia *et al*., 2017; Morse et al., 2024; Tresenrider *et al*., 2021). There is no expression of the canonical protein from the LUTI (hence they are “<u>u</u>ndecoded”) due to the presence of multiple uORFs at the 5’ end of the transcript. Transcript isoform switching between LUTIs and canonical transcripts occurs frequently in meiosis (Chen *et al*., 2017; Cheng *et al*., 2018). LUTIs also have key roles in the unfolded protein response (UPR) (Van Dalfsen *et al*., 2018), and similar two-transcript systems have functions in other areas of biology (Houseley et al., 2008; Martens et al., 2004). Many SUTs and XUTs also map 5’ of coding genes (van Dijk *et al*., 2011; Wery *et al*., 2016; Xu *et al*., 2009), and most *DCP2*-sensitive transcripts map proximal to inducible genes involved in metabolism (Geisler *et al*., 2012). This suggests they may also play a role in regulating expression of canonical transcripts.

Here, we use RNA-seq and a novel 40S ribosome profiling strategy to identify alternative translation events (mainly uORFs and iORFs) on transcripts associated with NMD-sensitive genes. We report a plausible rationale to account for most NMD-sensitive transcripts. Intriguingly, we find many NMD-sensitive, 5’-extended transcripts that are associated with inducible genes and function as LUTIs. We show that the NMD-sensitive *DAL5* LUTI has a functional role in repressing expression of the canonical *DAL5* transcript.

## RESULTS

### Detection of premature termination events that cause NMD

To identify genes that increased in abundance in the *upf1*Δ strain (i.e., NMD targets) we performed RNA-seq on BY4741 (WT) and *upf1*Δ *S. cerevisiae* strains followed by differential expression analysis on reads from main open reading frames (ORFs) with DESeq2 (Love et al., 2014). Because NMD has been reported to target genes that, to varying degrees, retain an intron (He *et al*., 1993), intron read counts were also analyzed (Figure S1A). Combining both analyses after duplicate removal (4 genes were found in both datasets, see Methods) yielded a high confidence set of 552 genes that were significantly changed and upregulated >2-fold in *upf1*Δ cells (Figure 1A and B, and Table S1 Sheet1).

**Figure 1.**
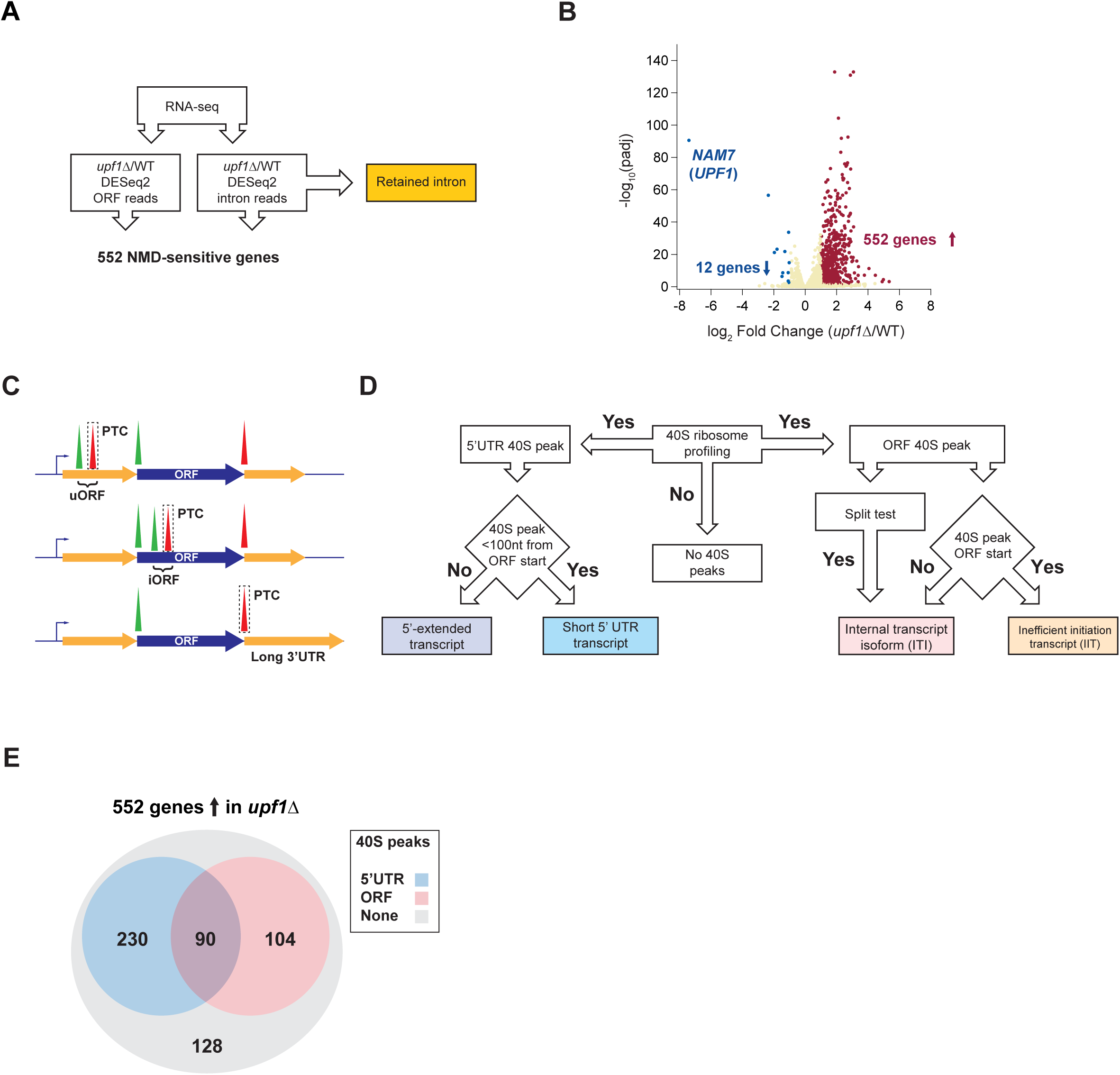
Strategy for identification of PTCs and 40S peak analysis. A) Pipeline for identification of NMD-sensitive genes, including genes with retained introns. DESeq2 was used to determine significantly changing RNA levels within ORFs and introns (552 total). Retained introns reported as a distinct category and other genes subjected to further analysis in D. B) Volcano plot shows cutoffs for significance and fold-change with each dot representing 1 gene. C) The three possible locations for PTCs: uORFs (top), iORFs (middle), main stop codon rendered a PTC by some other factor, such as long 3’UTR as shown (bottom). D) Pipeline for identification of 40S peak location to facilitate categorization of transcripts. E) Categorization of where 40S peaks were identified in NMD-sensitive genes.

We previously used 40S ribosome profiling to show that 40S subunits accumulate at start codons and that these peaks can be enhanced by reducing the level of 60S subunits via deletion of *RPL11B*, one of two genes that encode Rpl11p (uL5). We also showed that deletion of genes encoding 40S subunit recycling factors, *TMA64* and *TMA20,* results in a dramatic accumulation of 40S subunits at stop codons (Young et al., 2021). We reasoned that the peaks at start and stop codons could be utilized as markers to precisely identify novel translation events that would otherwise be missed by 80S ribosome profiling. To globally assess this association, we found that the 40S start or stop codon peak height of a given gene showed a strong correlation with the number of 80S ribosome footprints that mapped to the respective gene’s coding sequence (Figure S1B and C). Given the better correlation for start codon peaks, we focused on them for much of our analysis (see Methods). We therefore performed 80S ribosome profiling on WT and *upf1*Δ strains, and 40S ribosome profiling on *rpl11b*Δ and *tma64*Δ *tma20*Δ (abbreviated *64*Δ *20*Δ) in a *upf1*Δ strain background. Confirming this approach, we could detect a 40S stop codon peak at the previously identified PTC for the NMD target *RPL28* (*CYH2*) (Figure S1D).

Theoretically, there are three possible locations for premature termination events: after upstream open reading frames (uORFs) in the 5’ untranslated region (5’UTR) (uORFs; Figure 1C top), after internal out-of-frame ORFs within the main ORF (iORFs; Figure 1C middle), and “canonical” termination events that are rendered premature by some other feature of the transcript (Figure 1C bottom). We therefore developed a customized bioinformatic pipeline (Figure 1D) to identify putative premature termination events for the first two categories (Figure 1E) and offer examples and potential candidates for the third case where the canonical termination event is recognized as premature. As not every premature termination event triggers NMD, our approach focuses on the set of 552 NMD-sensitive genes we identified by RNA-seq and then offers a rationalization for the NMD sensitivity of each transcript.

### Classification of premature termination events in 5’UTRs

We detected 320 NMD-sensitive genes with 5’UTR translation events (Figure 1E) using 40S profiling peak score analysis (see Methods) (Table S1 Sheet2). Start codons were identified in all frames by extending the annotated 5’UTR upstream to capture any cases of alternative upstream TSSs (see Methods). As noted above, many yeast genes utilize multiple TSSs (i.e. LUTIs), and this results in the production of multiple transcripts per gene. Preliminary observations revealed two different classes of NMD-sensitive transcripts with uORFs: traditional annotated transcripts and a novel class of 5’-extended transcripts, often with multiple uORFs, similar to LUTIs (Figure 2A). Analysis of the distribution of yeast 5’UTR lengths based on a recent annotation of 5’UTR TSSs (Spealman et al. 2018) revealed that the average 5’UTR length is approximately 100 nt (Figure 2B). We therefore used 100 nt as a cut-off to define uORF-containing short 5’ UTR transcripts (Table S1 Sheet 3) vs 5’-extended transcripts (Figure 2C and Table S1 Sheet4).

**Figure 2.**
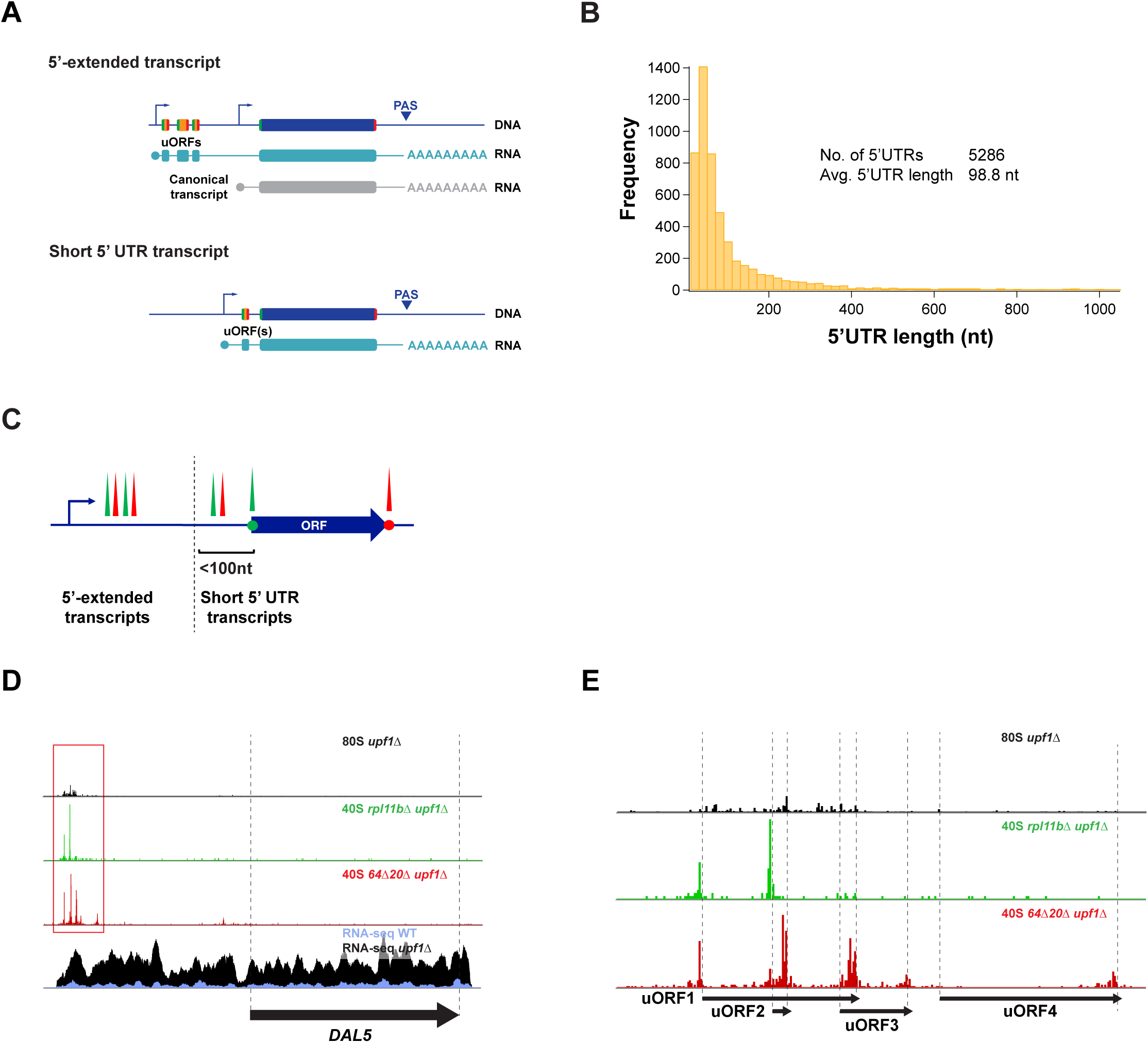
Categorization of 5’UTR 40S peak containing transcripts. A) Potential transcription architecture of yeast genes with uORFs: 5’-extended and canonical transcripts (top) or a single short transcript (bottom). Note the similarity of the architecture of the 5’-extended and canonical transcripts to the architecture of LUTI 2-transcript systems. B) Distribution of annotated most abundant 5’UTR lengths in yeast grown in glucose. C) Categorization of NMD-sensitive genes with 5’UTR 40S peaks into 5’-extended transcripts (184 genes) and short 5’UTR transcripts (136 genes) based on the average 5’ UTR length (100-nt cutoff). D) Sequencing reads for the *DAL5* gene in rich media (arrow corresponds to main ORF) show an example of a 5’-extended isoform. 80S ribosome profiling (top) reveals translation on uORFs. 40S profiling (middle tracks) shows locations of start and stop codons on uORFs. mRNA-Seq (bottom) shows expression of 5’extended transcript and its sensitivity to NMD. E) Close-up of D to reveal peaks on uORFs.

We identified the *DAL5* gene as an NMD-sensitive 5’-extended transcript (Figure 2D). The gene’s main ORF encodes allantoate permease and is repressed by nitrogen catabolite repression (NCR) under rich nitrogen growth conditions (Rai et al., 1988; ter Schure et al., 2000). The 5’-extended transcript includes a 5’UTR that measures approximately 1500 nt in length and includes 33 AUG codons that could serve as uORF start codons. We observed 40S start and stop codon peaks at the first four uORFs of the 5’-extended transcript (Figure 2E). Interestingly, these uORFs appear to inhibit expression of the main *DAL5* ORF based on the lack of 80S profiling reads (Figure 2D). In contrast, an example of an NMD-sensitive gene with a uORF that was classified as a short 5’ UTR transcript is *QDR3* (Figure S1E).

### Classification of premature termination events in coding sequences

We detected 194 NMD-sensitive genes with internal translation events within the ORF in a frame different from the coding sequence (Figure 1E) using 40S profiling peak score analysis (see Methods) (Table S1 Sheet5).

There are several possible explanations for the observed translation events within main ORFs: (1) The 48S pre-initiation complex (PIC) does not recognize the annotated start codon. This could be due either to poor start codon context (leaky scanning) or a short transcript leader that renders the PIC unable to scan until downstream of the annotated start codon. Both alternatives should result in initiation at the next available start codon within the main ORF. We call these collectively inefficient initiation transcripts (IITs; Figure 3A top). (2) The TSS for the transcript is within the main ORF, resulting in a 5’ truncated transcript. This internal transcript isoform (ITI) lacks the canonical ORF start codon so translation must initiate at an internal AUG (Figure 3A bottom).

**Figure 3.**
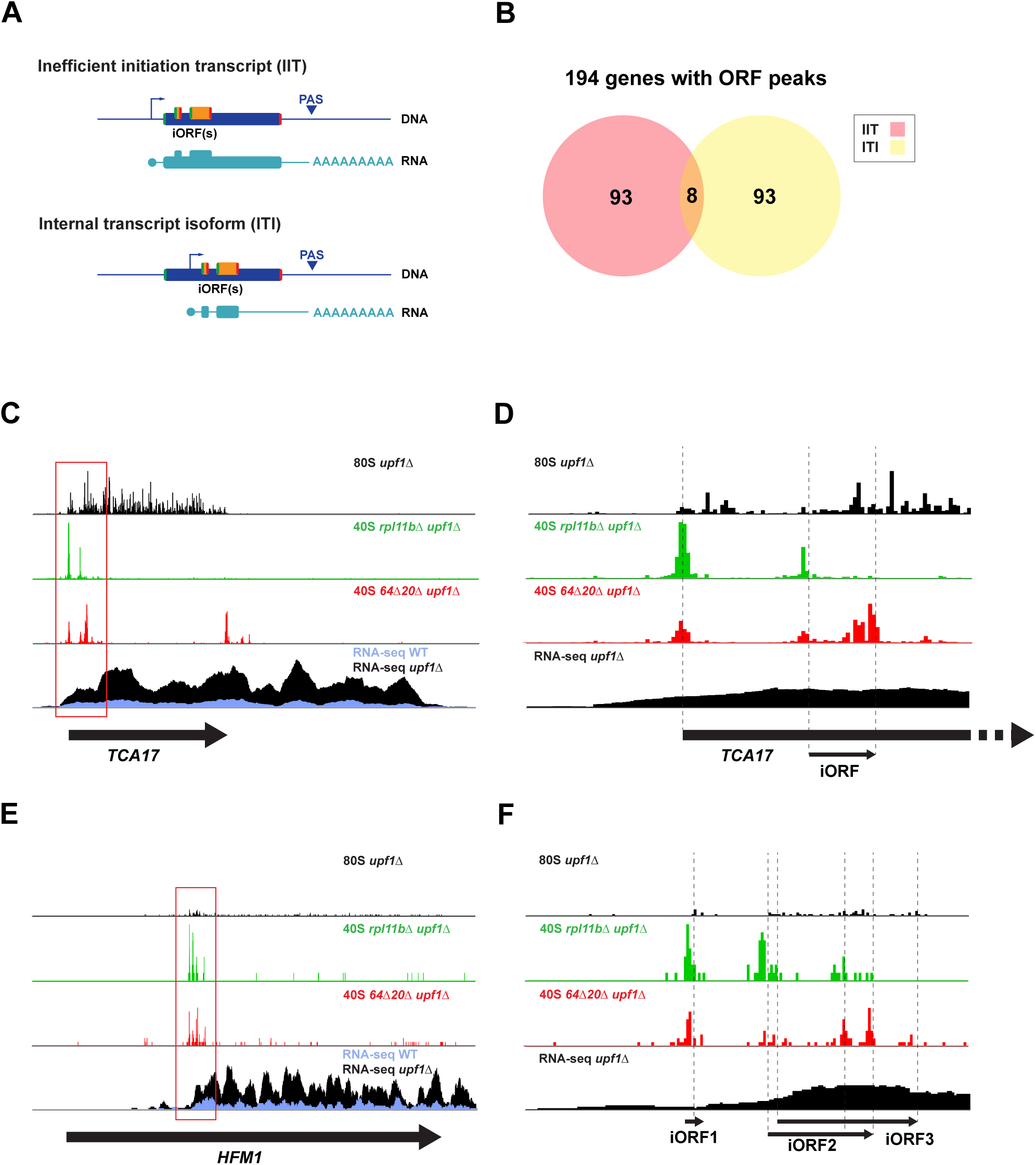
Categorization of ORF 40S peak containing transcripts. A) Potential transcription architecture of yeast genes with iORFs: inefficient initiation transcript (ITI) of a canonical isoform due to leaky scanning or (as shown) or short transcript leader (top) or internal transcript isoform (ITI) where transcript starts within the main ORF (bottom). B) Categorization of NMD-sensitive genes with main ORF 40S peaks into inefficient initiation transcripts (IITs) or internal transcript isoforms (ITIs) based on presence or absence of a 40S main ORF start codon peak and the split-gene test. C) Sequencing reads for the *TCA17* gene (arrow corresponds to main ORF) show an example of an IIT. 80S ribosome profiling (top) fails to show iORF translation. 40S profiling (middle tracks) shows locations of start and stop codons for the iORF. mRNA-Seq (bottom) shows expression of canonical transcript and its sensitivity to NMD. D) Close-up view of C to show individual peaks on translated iORF. E) Sequencing reads for the *HFM1* gene (arrow corresponds to main ORF) show an example of an ITI. 80S ribosome profiling (top) shows translation of iORFs. 40S profiling (middle tracks) shows locations of start and stop codons on iORFs. mRNA-Seq (bottom) shows expression of ITI and its sensitivity to NMD. F) Close-up view of E to show individual peaks on translated iORFs.

We divided NMD-sensitive genes with 40S ORF peaks into IITs, where the start codon is present though used inefficiently, and ITIs, where the canonical start codon is absent, using several bioinformatic criteria. A simple way to separate IITs and ITIs is based on the presence or absence of a 40S peak at the start codon of the main ORF (Figure S2A). NMD-sensitive genes with 40S start peaks were categorized as IITs (101 genes) (Figure 3B; Table S1 Sheet6), whereas genes without were categorized as ITIs (93 genes) (Table S1 Sheet7). Another bioinformatic method to identify ITIs based on *upf1*Δ RNA-seq data was developed (see Methods). This split-count test (Figure S2B) compared RNA-Seq read density in the first and last 100 nt of the main ORF. Those with a more than 2-fold higher read count in the final 100 nt of the ORF relative to the first 100 nt were identified as ITIs (49 genes) (Table S1 Sheet8). Comparison of the two tests showed strong overlap (Figure S2C), so the lists were combined to yield a total of 101 ITIs (Figure 3B; Table S1 Sheet9). The *TCA17* gene expresses an IIT that appears to inefficiently initiate translation at the main start codon, possibly due to a short (20 nt) 5’UTR (Figure 3C), and often initiates translation on a 7-aa iORF (Figure 3D). The *TCA17* gene reveals the power of the 40S profiling technique as the internal translation event cannot be detected in the 80S ribosome profiling track. The *HFM1* gene expresses an ITI (Figure 3E) that exhibits 40S start and stop codon peaks at three out-of-frame iORFs within the ORF (Figure 3F).

### Overlap and additional causes for peaks

We noted that many of the 90 genes (Figure 1E) that had both 5’UTR and ORF peak(s) appear to express two NMD-sensitive transcript isoforms, such as *BDS1* (Figure S2D). In addition, our 40S profiling method identified programmed ribosomal frameshift sites. Of 3 known (Farabaugh et al., 2006; Palanimurugan et al., 2004) sites, *OAZ1*, *EST3*, and *ABP140*, only *EST3* shows significant NMD-sensitivity (Figure S2E). In total, we found a translation event that likely results in premature termination for 77% of the 552 NMD-sensitive genes we identified by 40S peak analysis (Figure 1E).

### Confirming the role of 5’ uORFs in targeting DAL5 to NMD

To confirm the ability of uORFs to trigger NMD on 5’-extended transcripts, we developed a system (based on the YCplac33 sc *URA3* plasmid) for studying the 5’-extended transcript of *DAL5* (Figure 2D and 2E and Methods). Northern blots of RNA samples from YCplac33-*DAL5* cells recapitulated what we observed in our RNA-seq data with the 5’-extended *DAL5* transcript being stabilized in the *upf1*Δ background strain (Figure 4A). Having successfully established a system for studying the 5’-extended transcript of *DAL5*, we then mutated the 33 AUG start codons in the 5’-extended transcript upstream of the YCplac33-*DAL5* plasmid to AAA to test if these uORFs target the transcript for NMD (Figure S3A and B). Mutation of the 5’ uORF AUGs resulted in stabilization of the *DAL5* 5’-extended transcript in *UPF1*^+^ (WT) cells (Figure 4B), proving that the uORFs are responsible for the transcript’s NMD sensitivity. The level of stabilization is equivalent to that observed in the *upf1*Δ strain, suggesting that these uORF termination events account for all the NMD sensitivity. To test the generality of this result, we repeated the experiment with another NMD-sensitive gene with a long transcript, *DAL7* (Figure 4C). While the *DAL7* 5’UTR does meet the 100-nt cutoff to be classified as a 5’-extended transcript, experiments below suggest it is part of a two-transcript system. A similar plasmid-based system was established to mutate its 3 AUG start codons (Figure S3C and D). As with *DAL5*, these mutations resulted in the stabilization of *DAL7* (Figure 4D). These results show that PTCs that we identified with 40S ribosome profiling target these transcripts to NMD.

**Figure 4.**
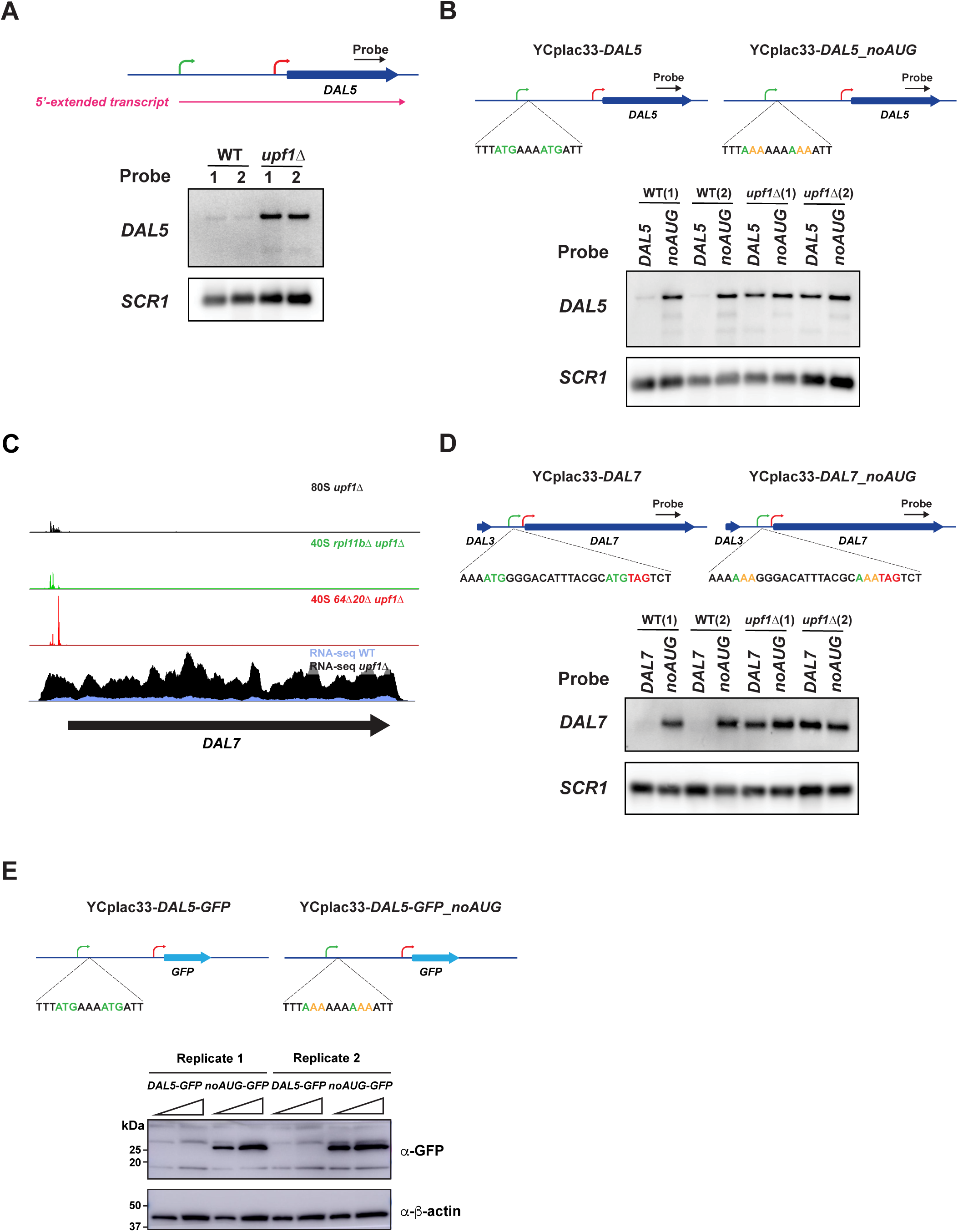
uORFs in the 5’-extended transcripts of *DAL5* and *DAL7* are responsible for NMD sensitivity. A) Northern analysis of the *DAL5* 5’-extended transcript expressed from the YCplac33-*DAL5* plasmid confirms NMD sensitivity. B) Northern analysis of the *DAL5* 5’-extended transcript expressed from YCplac33-*DAL5* and YCplac33-*DAL5_noAUG* where all uORF AUGs are mutated to AAA shows that the *DAL5* uORFs confer NMD sensitivity. C) Sequencing reads for the *DAL7* gene in rich media (arrow corresponds to main ORF). 80S ribosome profiling (top) reveals translation on uORFs. 40S profiling (middle tracks) shows locations of start and stop codons on uORFs. mRNA-Seq (bottom) shows expression of the long transcript and its sensitivity to NMD. D) Northern analysis of *DAL7* long transcript expressed from YCplac33-*DAL7* and YCplac33-*DAL7_noAUG* where all uORF AUGs are mutated to AAA shows uORFs fully confer NMD sensitivity. E) Western analysis of the YCplac33-*DAL5*-*GFP* and YCplac33-*DAL5*-*GFP*_*noAUG* reporters show that loss of all uORFs in the *DAL5* 5’extended transcript leader leads to translation of the main ORF. Second lane for each experiment is 2X loaded. In blots above, *SCR1* and β-actin serve as loading controls. Two replicates are shown for each condition.

Given the length of the *DAL5* 5’-extended transcript’s 5’UTR (∼1.5 kb), we wondered whether, in the absence of 5’ uORFs, the 48S PIC would be able to scan the entire 5’UTR and initiate translation at the main ORF start codon. We therefore constructed versions of YCplac33-*DAL5* and YCplac33-*DAL5_noAUG* where the *DAL5* main ORF was replaced with the coding sequence for *GFP*. We found that, in the absence of 5’ uORFs, GFP was expressed (Figure 4E), demonstrating that the 5’ uORFs in the 5’-extended transcript are critical for preventing expression of the Dal5 protein.

### Potential role of long 3’UTRs in stimulating NMD

We found that 128 genes (including those with retained introns) that increased in abundance in the *upf1*Δ strain lack 40S peaks (Figure 1E). In many cases, therefore, we are left to conclude that it is the main stop codon that is recognized as a PTC (Figure 1C bottom). While multiple factors could be responsible for this effect, previous studies have shown that inefficient termination at a canonical stop codon can be driven by a long 3’UTR (Kebaara and Atkin, 2009; Muhlrad and Parker, 1999; Peccarelli *et al*., 2016). For example, the *COX19* gene has been reported to be targeted to NMD and 3’UTR swapping experiments showed that the long 3’UTR is responsible (Peccarelli *et al*., 2016). We therefore examined the distribution of 3’UTR lengths (Figure S4A left) and chose the average plus two standard deviations (>395 nt) to generate a list of 75 genes with “long” 3’UTRs. While we found this 3’UTR length was, on average, more sensitive to NMD (Figure S4A right), it is not robust. We therefore utilized a process of elimination to find the best cases where a transcript’s 3’UTR length is a likely trigger NMD and offer a resource for future study (Figure S4B).

First, we noted that in pioneering work, (Kebaara and Atkin, 2009) bioinformatically identified 56 genes with 3’UTRs >350 nt as strong candidates where 3’UTR length would drive NMD (Kebaara and Atkin, 2009) - Supplemental Table 1 therein). We found that 19 of these genes were significantly NMD-sensitive and exhibited 3’UTRs with a predicted length similar to that observed in our RNA-seq data (Table S1 Sheet10). Second, based on our analysis (Figure S4A), we used the 395 nt cutoff to add 27 NMD-sensitive genes where a long 3’UTR could be the driver of NMD (Table S1 Sheet11). Consistent with the main stop codon being recognized as a PTC, the main ORF exhibited an increase in 80S ribosome profiling footprints in the *upf1*Δ strain in all these cases.

Next, we considered the observation that “bicistronic” mRNAs had been previously suggested as a type of NMD substrate (He *et al*., 2003). The use of the term bicistronic can be misleading; it is unlikely that these transcripts produce two protein products from a single mRNA. We therefore suggest that a more accurate term is *pseudo-bicistronic transcripts*. Pseudo-bicistronic transcripts are potential NMD targets since translation termination at the stop codon of the first ORF could be recognized as a PTC with the second ORF effectively acting as a long 3’UTR. In many cases, a second transcript covering the second ORF in the pair is also expressed, thus enabling production of both protein products (Figure S4C).

A study using TIF-Seq identified 279 bicistronic transcripts in yeast (Pelechano *et al*., 2013) - Supplementary data 10 therein). After manual editing for false positives (see Methods), we found that 50 bicistronic transcripts where either one or both genes (68 total) were NMD targets in our RNA-Seq data (Table S1 Sheet12). Anecdotally, we noticed an additional 13 genes involved in pseudo-bicistronic pairs that did not meet the threshold defined as bicistronic transcripts in the TIF-seq study (Pelechano *et al*., 2013). These genes were part of 9 pseudo-bicistronic transcripts (Table S1 Sheet13). *SLO1-ISC10* is an example of a gene pair expressing an NMD-sensitive pseudo-bicistronic transcript originally identified by TIF-seq (Figure S4D). Our data show that only *SLO1* is translated, making it likely that its canonical stop codon acts as a PTC. In another example, (Kebaara and Atkin, 2009) determined that the *MAK31* gene is part of two isoforms, the longer of which is NMD-sensitive and constitutes a pseudo-bicistronic transcript extending over the *PET18* ORF (Figure S4E). Our data again shows that only *MAK31* is translated with its canonical stop codon acting as a likely PTC.

Lastly, we considered pseudogenes as another case that effectively acts as a long 3’UTR. We define a pseudogene here as a tandem gene pair separated by a stop codon or other mutation that is not present in closely related *Saccharomyces* species (see Methods). We identified 22 NMD-sensitive genes that are paired into 11 likely pseudogene transcripts (Table S1 Sheet14). As an example, the *NIT1-YIL165C* pseudogene encodes a nitrilase with a single premature stop codon (Figure S4F).

Taken together, the above analysis and observations argue that long 3’UTRs could be a driver of NMD for many genes, including many that may be targeted for additional reasons due to 40S peaks we identified above. Of the 128 genes with no 40S peaks, we suggest that long 3’UTRs, where the canonical stop codon acts as a premature stop codon, and pseudogenes may explain NMD-sensitivity for an additional 60 genes. With the 424 genes with 40S peaks, and 10 genes with retained introns our analysis offers a rationalization of NMD sensitivity for 494 of 552 genes or 89%.

### NMD-sensitive LUTIs regulate metabolism

The NMD-sensitive 5’-extended transcripts identified here resemble LUTIs shown previously to play a role in a two-transcript gene regulatory system (Figure 2A top) (Chen *et al*., 2017; Cheng *et al*., 2018; Chia *et al*., 2017; Tresenrider *et al*., 2021). Genes utilizing this system have two promoters: a proximal promoter close to the main ORF from which a canonical (or induced) transcript isoform is expressed and a 5’ distal promoter from which the LUTI is expressed. Transcription of the canonical transcript from the proximal promoter results in translation of the main ORF protein (uORFs block main ORF translation on LUTI transcripts). Transcription factors have been proposed to bind to both promoters and respond to changing environmental conditions or developmental programs, though the magnitude of change (particularly for the LUTI transcript, which does not always change) varies considerably between genes (Tresenrider *et al*., 2021). The key function of the LUTI is that its transcription represses the downstream proximal promoter via repressive chromatin marks (Chia *et al*., 2017; Morse *et al*., 2024; Tresenrider *et al*., 2021). We compared the list of genes with putative LUTIs associated with meiosis from (Cheng *et al*., 2018) [Table S2 therein] with our list of 320 NMD-sensitive genes with 5’ UTR 40S peaks and found 84 in common. This overlap suggests that NMD-sensitive 5’-extended transcripts have functional roles as LUTIs. To better assess whether the NMD-sensitive 5’-extended transcripts we observed in rich media are LUTIs, we asked whether a second, canonical transcript could be induced under alternate growth conditions. Similar to previous studies of NMD-targets (He *et al*., 2003; Johansson *et al*., 2007), many genes in our list are involved in nitrogen catabolism and thiamine biosynthesis. We therefore grew WT and *upf1*Δ cells under either poor nitrogen or no thiamine growth conditions and performed RNA-seq.

We identified 48 genes that were >5-fold upregulated in poor nitrogen (Table S1 Sheet15). We then compared changes in expression due to poor nitrogen with changes in expression due to loss of *UPF1* (Figure 5A). We identified a group of genes with NMD-sensitive long transcripts that showed increased expression in poor nitrogen (upper right quadrant), suggestive of a LUTI architecture (Figure 5A). To further investigate if these transcripts were LUTIs, we focused on the *DAL5* and *DAL7* genes. Both expressed shorter, canonical transcripts that were not sensitive to NMD under poor nitrogen conditions in addition to NMD-sensitive long transcripts that were constitutively expressed under rich and poor conditions (Figure 5B and S5A). To confirm these observations, we performed northern blots under poor nitrogen conditions (Figure 5C and S5B). We observed similar stability patterns for the long (candidate LUTI) transcripts as we did under rich nitrogen growth conditions (Figure 4B and D, i.e., loss of uORFs stabilized the extended transcripts). We also observed distinct shorter, induced *DAL5* and *DAL7* transcripts that were not subject to NMD, confirming that at least two transcript isoforms are expressed from these genes. Interestingly, we noted that low-nitrogen conditions revealed that the *NIT1-YIL165C* pseudogene that we described above (Figure S4F) expressed two transcripts that were both sensitive to NMD (Figure S5C). A shorter transcript was induced under low nitrogen and a longer transcript, likely functions as a LUTI repressing it. The shorter induced pseudogene transcript contains a PTC that also targets it for NMD (Figure S5E compare WT+ proline to *upf1*Δ + proline). These findings establish that 5’-extended transcripts are associated with inducible transcripts involved in NCR and are strong candidates for LUTIs.

**Figure 5.**
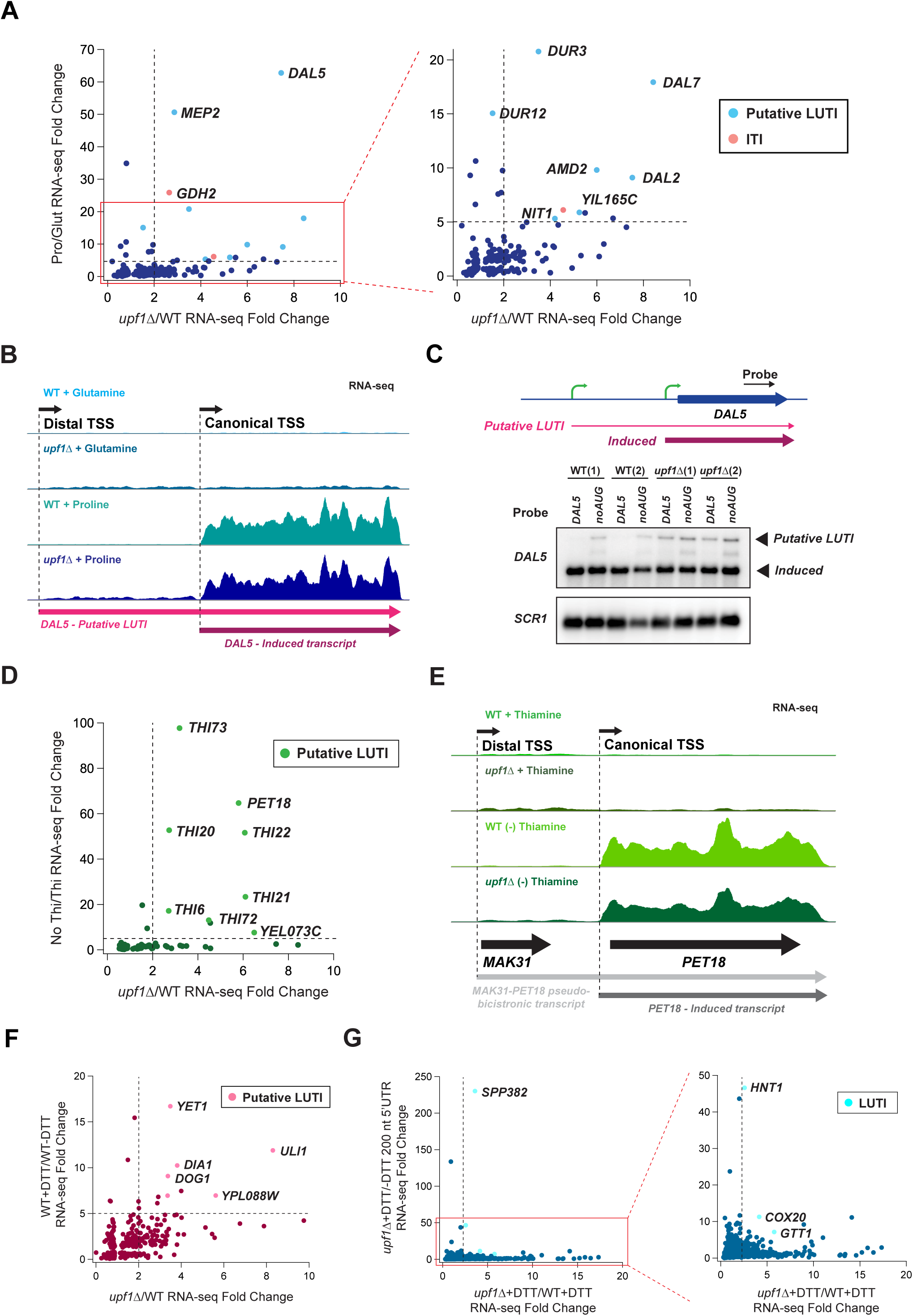
Identification of regulated transcript pairs that include an NMD-sensitive 5’-extended (LUTI) transcript and downstream (canonical) transcript. A) Comparison of differential expression in mRNA-Seq data for poor vs rich nitrogen (y-axis) vs loss of *UPF1* (x-axis). Genes most affected by both conditions (upper right quadrant of zoom-up) are enriched for putative LUTIs. B) mRNA-Seq reads mapped to the *DAL5* gene in order from top to bottom: WT in nitrogen-rich media, *upf1*Δ in nitrogen-rich media, WT in nitrogen-poor media, *upf1*Δ in nitrogen-poor media. Black arrow on top show approximate TSSs. Colored arrows at bottom show transcripts. Data reveal that the NMD-sensitive, putative LUTI is present under both media conditions and the existence of a shorter canonical NMD-insensitive transcript that is induced under poor nitrogen. C) Northern analysis confirms expression of both a *DAL5* putative LUTI and a canonical induced transcript under poor nitrogen conditions. The 5’ uORFs confer NMD sensitivity to only the putative LUTI. *SCR1* serves as a loading control. Two replicates are shown for each condition. D) Comparison of differential expression in mRNA-Seq data for loss of thiamine (y-axis) vs loss of *UPF1* (x-axis). The genes most affected by both conditions are enriched for putative LUTIs. E) mRNA-Seq reads mapped to the *MAK31-PET18* pseudo-bicistronic gene pair in order from top to bottom: WT in media with thiamine, *upf1*Δ in media with thiamine, WT in media without thiamine, *upf1*Δ in media without thiamine. Black arrows on top show approximate TSSs. Arrows at bottom show transcripts. Data reveal an NMD-sensitive, pseudo-bicistronic transcript that is present under both media conditions and an NMD-insensitive induced *PET18* transcript that is only present in the absence of thiamine. F) Comparison of differential expression in mRNA-Seq data for +DTT vs −DTT (y-axis) vs loss of *UPF1* (x-axis). The genes most affected by both conditions are enriched in putative LUTIs, such as *ULI1*. G) Comparison of differential expression in mRNA-Seq data over the region 200 nt upstream of the start codon for +DTT vs −DTT (y-axis) vs loss of *UPF1* (x-axis). This analysis reveals cases where NMD-sensitive LUTIs may be turned on by DTT. For A, D, F, and G, included differentially expressed genes are those that have FDR<0.01 in both experiments.

We identified 23 genes that were >5-fold upregulated in the absence of thiamine (Table S1 Sheet16). As above with poor nitrogen, we then compared changes in expression due to loss of thiamine with changes in expression due to loss of *UPF1* (Figure 5D). Similarly, we again identified a group of genes with NMD-sensitive long transcripts that showed increased expression in the absence of thiamine (upper right quadrant), suggestive of a LUTI architecture (Figure 5D). The *PET18* gene, which was discussed above as part of an NMD-sensitive *MAK31-PET18* pseudo-bicistronic transcript (Figure S4E), and the *THI22* gene, which expresses a 5’-extended transcript, expressed canonical transcripts in the absence of thiamine (Figure 5E and S5D). These results were previously confirmed by northern blotting (Johansson *et al*., 2007) [Figure 3A therein]. Similar to *MAK31-PET18* and *THI22*, *THI6* appears to be a two-transcript system with a short, induced transcript expressed in the absence of thiamine, but unlike the others, the longer transcript appears to turn off in the absence of thiamine (Figure S5E). These examples demonstrate how 5’-extended transcripts in the thiamine sensing pathway likely function as LUTIs.

Many LUTIs are known to function in the UPR (Van Dalfsen et al. 2018). To explore whether any of the 5’-extended transcripts that we found to be sensitive to NMD were involved in UPR regulation, we treated WT and *upf1*Δ cells with 5 mM DTT to induce ER stress and performed RNA-seq (Figure 5F). As with the stress conditions above, we noted several genes that were sensitive to both NMD and DTT, including *ULI1*. *ULI1* expresses a canonical transcript that is activated by DTT and a 5’extended transcript appears to behave as a LUTI. To identify LUTIs that are induced by DTT, but masked by NMD, we looked for increases in RNA-seq for the 200 nt upstream of the ORF in *upf1*Δ cells treated with DTT versus untreated (Figure 5G, see Methods). This analysis revealed previously identified LUTIs (*GTT1*, *COX20*, *HNT1*) that we show here are NMD-sensitive and identify a candidate LUTI for *SPP382*.

### Transcriptional regulation of DAL5 and other NMD-sensitive LUTIs

To more firmly establish that the 5’-extended transcript of *DAL5* functioned as a LUTI, we next investigated whether it could repress the transcription of its corresponding downstream canonical transcript. To test this, we asked whether prematurely terminating the *DAL5* 5’-extended transcript derepressed the canonical (inducible) *DAL5* promotor and resulted in increased expression of the canonical transcript. To terminate the *DAL5* 5’-extended transcript, we inserted a truncated version of the *SNR13* transcription terminator sequence at several locations within the 5’-extended region of the *DAL5* gene (Porrua et al., 2012) and prior to the canonical *DAL5* promoter (Figure 6A). We transformed these plasmids into a *upf1*Δ strain (that also lacked the chromosomal region corresponding to the *DAL5* insert in YCplac33-*DAL5*, *dal5-ext*Δ) so that the 5’-extended transcript would be stabilized. We grew cells in rich nitrogen conditions (YPD), where the canonical *DAL5* transcript should be repressed. Northern blotting showed that truncation of the 5’-extended *DAL5* transcript resulted in derepression of a proximal *DAL5* promoter (Figure 6B). This result shows that the *DAL5* 5’-extended transcript functions as a LUTI transcript by repressing the downstream canonical (induced) transcript. We repeated this experiment under poor nitrogen conditions, where the canonical transcript is induced, and observed that truncation of the *DAL5* LUTI resulted in an increase in the amount of induced transcript (Figure S6A). The role of the 5’-extended (LUTI) transcript would therefore appear to be functionally important in both silencing the canonical transcript under non-inducing conditions and limiting its expression under inducing conditions. We note that a similar experiment was previously conducted on *DCI1*, a gene encoding an enzyme involved in fatty acid oxidation, which we identified as expressing an NMD-sensitive 5’-extended transcript. Previously, (Kim et al., 2012) showed that repression of a shorter induced (canonical) mRNA was dependent on expression of the longer RNA. The example of *DCI1* further establishes 5’-extended NMD-sensitive transcripts serve as LUTIs.

**Figure 6.**
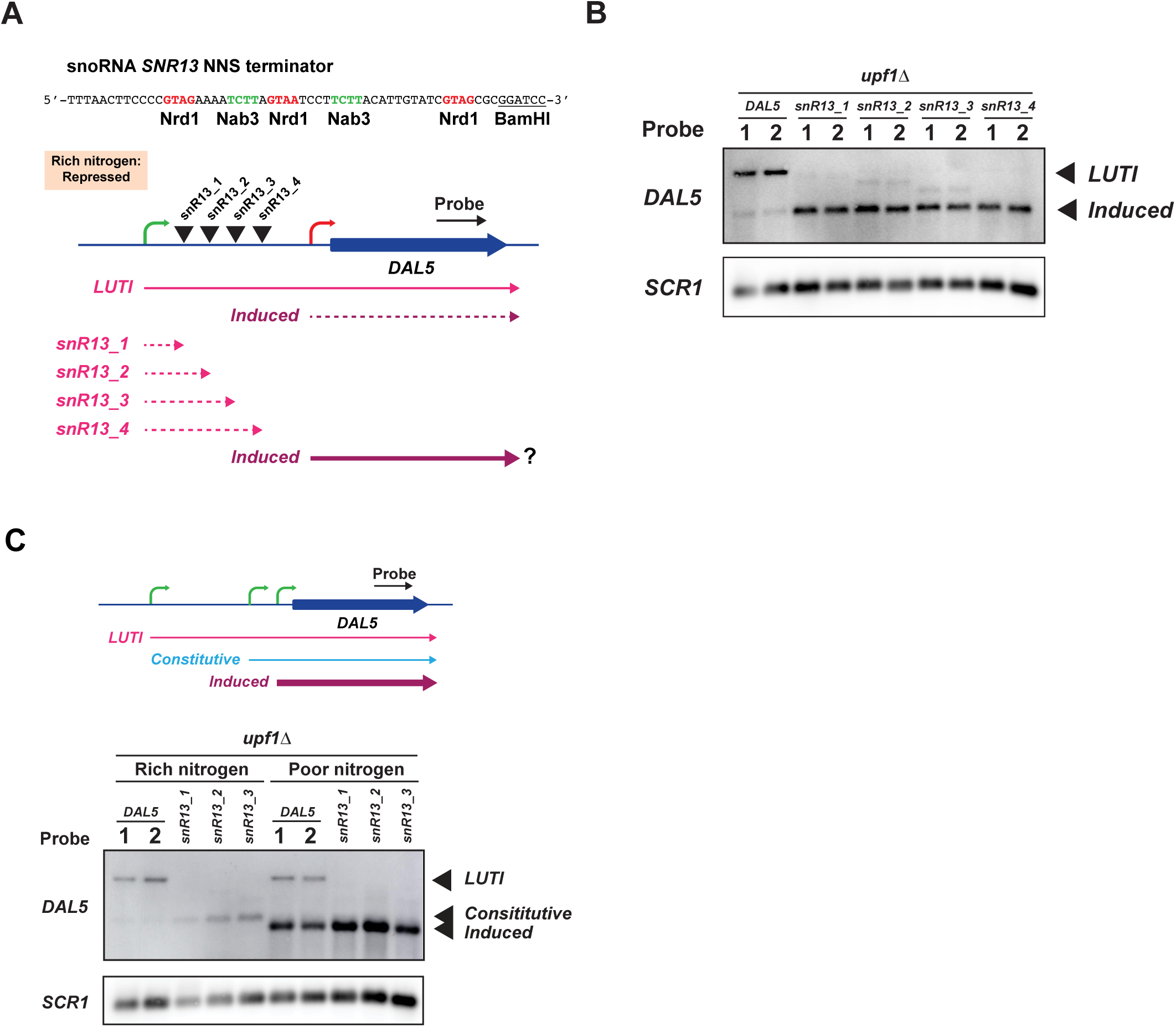
Transcription of the *DAL5* 5’-extended LUTI represses two downstream promoters. A) Design of experiment to test role of *DAL5* LUTI transcription in repressing downstream transcription. The truncated snoRNA *SNR13* NNS terminator was inserted at four locations within the 5’-extended transcript of the YCplac33-*DAL5* plasmid. B) Northern analysis of *DAL5* shows premature termination of the *DAL5* LUTI transcript derepresses a downstream transcript under conditions of rich nitrogen. *SCR1* serves as a loading control. Two replicates are shown for each condition. C) Northern analysis of *DAL5* shows premature termination of the LUTI transcript derepresses two downstream transcripts as evidenced by the appearance of a constitutive transcript under rich nitrogen and the induced transcript under poor nitrogen. *SCR1* serves as a loading control.

Transcription of LUTIs represses the canonical promoter by co-transcriptional recruitment of chromatin modifying enzymes that establish a repressive chromatin state (Chia *et al*., 2017; Morse *et al*., 2024; Tresenrider *et al*., 2021). In particular, the Set1 and Set2 histone methyltransferases deposit H3K4me2 and H3K36me3 to recruit the histone deacetylase complexes Set3C and Rpd3S (Chia *et al*., 2017; Tresenrider *et al*., 2021). LUTI-based regulation is therefore dependent on this machinery. We noted that deletion of *SET3* (encoding the core subunit of the Set3C complex) results in derepression of canonical transcripts for *DCI1* and *DUR3* (Kim *et al*., 2012); shown in Figure 3A therein), and deletion of *SET2* results in derepression of *AAD10* (Kim et al., 2016); shown in Figure 2A therein), *YNR068C*, *DAL5*, *SSA3*, and *ZRT1* (Kim *et al*., 2016); shown in Figure S2A therein). We identified these genes as expressing NMD-sensitive 5’-extended transcripts, pseudo-bicistronic transcripts, or a split pseudogene. This adds support that these genes express LUTIs that are sensitive to NMD. In further support, CAGE-seq data from *set1*Δ *set2*Δ cells confirmed the derepression of the canonical transcript for *DAL5* (Figure S6B) and a transcript that only included the second ORF in the pairs *YJR154W-AAD10 and AAR2-SSA3* (Figure S6C; (Poonia et al., 2025).

Running the *DAL5* NNS terminator samples grown in rich and poor nitrogen media on the same northern blot revealed a surprising result (Figure 6C). The *DAL5* LUTI transcript was the same size in both rich and poor nitrogen conditions, however, the shorter (derepressed) transcript was not. This suggests the *DAL5* LUTI suppresses two transcripts: the longer of the two transcripts is derepressed in rich nitrogen and we shall therefore refer to it as the *DAL5* constitutive transcript and the shorter is derepressed by poor nitrogen and we shall therefore refer to it as the *DAL5* induced (or canonical) transcript. CAGE-seq data (Poonia *et al*., 2025) for yeast grown under rich nitrogen conditions suggest a start site for the *DAL5* constitutive transcript showing evidence for 4 uORFs prior to the annotated *DAL5* start codon that likely prevent protein production and sensitize the transcript to NMD. We confirmed that the *DAL5* constitutive transcript is targeted to NMD by repeating the *DAL5* NNS terminator experiment in WT (*UPF1*^+^) cells (Figure S6D). Based on RNA-Seq data, the induced canonical *DAL5* transcript likely contains no uORFs, explaining its lack of sensitivity to NMD (Figure 5B and C), as expected for a transcript needed to produce Dal5 protein. These results suggest the *DAL5* LUTI serves the dual role of suppressing the constitutive (rich nitrogen conditions) and induced (poor nitrogen conditions) transcripts.

### The novel DAL5 regulatory system may be important for function

Beyond regulation under steady state, it is conceivable that the *DAL5* LUTI plays a role in regulating the kinetics of Dal5 protein production under changing conditions of environmental nitrogen. To test this, we grew *upf1*Δ *dal5_ext*Δ cells carrying WT (YCplac33-*DAL5*) and an NNS termination mutant (YCplac33-*DAL5-SNR13_1*) in rich nitrogen and then transferred the cells to poor nitrogen and followed the induction of the induced *DAL5* transcript (Figure 7A). As demonstrated above, truncation of the *DAL5* LUTI transcript resulted in a higher endpoint of induction (Figure 7A and S6E), and in addition, the induction occurred at a higher rate. We also performed repression experiments where we grew cells in poor nitrogen to log phase and then transferred the cells to rich nitrogen (Figure 7B). Truncation of the *DAL5* LUTI resulted in a decreased rate of repression of not only the induced *DAL5* transcript but also the constitutive *DAL5* transcript (Figure 7B). These results show that the *DAL5* LUTI is important for the NCR response by tempering induction in poor nitrogen and accelerating shut-off upon return to rich nitrogen conditions.

**Figure 7.**
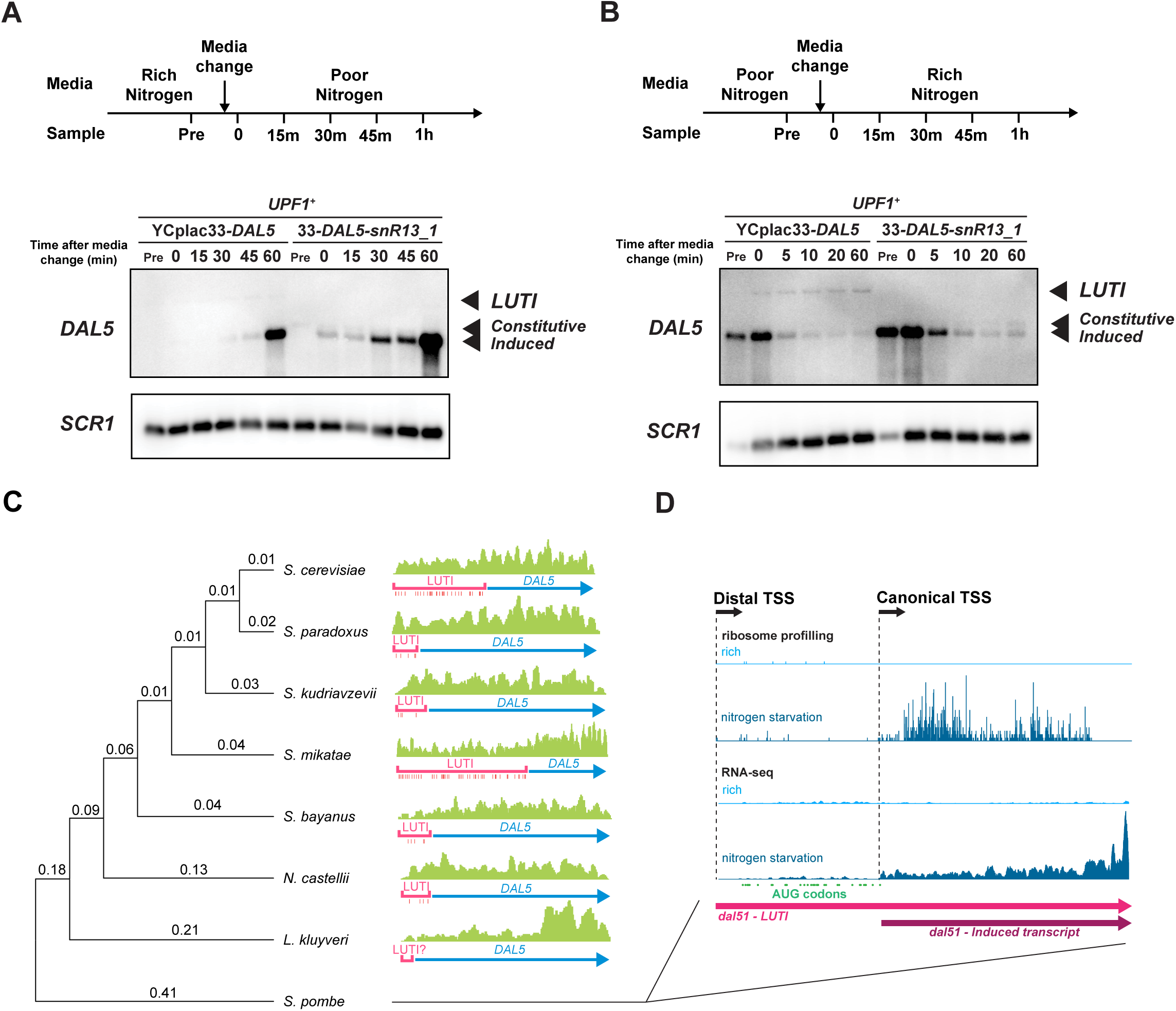
The *DAL5* LUTI regulates the kinetics of expression of the induced *DAL5* transcript. A) Northern analysis of *DAL5* shows derepression of the induced transcript after media change to poor nitrogen is more rapid, in addition to resulting in a higher endpoint, in the absence of the LUTI transcript. B) Northern analysis of *DAL5* shows repression of the induced transcript after media change to rich nitrogen is more rapid in the absence of the LUTI transcript. C) Phylogenetic tree (left) shows the evolutionary relationships among DAL5 proteins from related yeasts, with the numbers on the branches indicating the estimated number of amino acid substitutions per site relative to the parent node. RNA-seq data for the *DAL5* ortholog of each species is shown on the right. The blue arrow corresponds to the main ORF of the gene and the pink arrow was drawn to highlight the transcript expressed under rich conditions, based on the level of RNA-Seq reads. The 5’UTR of the *DAL5* ortholog is long in most species and contains multiple uORFs (vertical bars indicate AUG positions), consistent with the presence of a LUTI. Data from *L. kluyveri* was ambiguous on whether there was a long 5’UTR with uORFs that would constitute a LUTI. Data for *S. pombe* from a more expansive set of data is shown in D. D) Sequencing reads for the *dal51* (*DAL5* ortholog in *S. pombe*) gene show conservation of the LUTI architecture. Data from WT *S. pombe* cells. 80S ribosome profiling (top) reveals translation of main ORF under conditions of nitrogen starvation. mRNA-Seq (bottom) shows expression of the LUTI under rich nitrogen conditions and expression of the canonical transcript under poor nitrogen conditions. AUG start codons of potential uORFs indicated (green dots). LUTI annotation taken from PomBase (Rutherford et al., 2024) and canonical transcript estimated based on change in RNA-Seq level. *SCR1* serves as a loading control in the above blots.

If these biological functions of the *DAL5* LUTI system are important, it is likely they are conserved between species. We therefore investigated previously published mRNA-Seq data in related species (Blevins et al., 2019). We found in nearly all cases evidence for a transcript that extended far upstream of the main ORF and included many uORFs, which are hallmarks of LUTIs (Figure 7C). We also examined ribosome profiling and mRNA-Seq from a previous study (Duncan and Mata, 2014) in a more distant species, *S. pombe,* in rich and poor nitrogen conditions. As in our data from S*. cerevisiae* (Figures 2D and 5B), we found long and short isoforms of the transcript, with the longer isoform including translated uORFs and the shorter isoform being induced by poor nitrogen (Figure 7D). The conservation of these features offers strong evidence of the LUTI architecture being conserved over considerable evolutionary distance.

## DISCUSSION

An ongoing challenge is to fully catalog the transcripts that NMD targets and, more broadly, understand their functional significance in the cell. Our study has addressed this with a novel experimental approach for identifying cryptic translation events for the majority of the 552 genes that we found to be targeted by NMD in *S. cerevisiae*. Our method utilizes 40S ribosome profiling, which clearly reveals where ribosomes initiate and terminate translation on cryptic ORFs that are often invisible to conventional 80S ribosome profiling (for example, *TCA17*, Figure 3C). Using this, we offer at least one candidate PTC for the vast majority of NMD targets (Table S2). In cases where 40S profiling did not reveal a candidate PTC, we utilized a qualitative approach to propose possibilities, such as long 3’UTRs (Figure S4B).

Related to the question of what determines NMD sensitivity is what is the functional role of NMD? While it is clear NMD can degrade transcripts where PTCs are included as errors, our analysis shows NMD also plays a major role in regulating mRNA levels. In some cases, these transcripts require degradation because they lack obvious function (i.e. ITIs, pseudogenes, and some pseudo-bicistronic transcripts) but in others it appears the transcript is functional and NMD controls its level (i.e. LUTIs, IITs, long 3’UTRs, and some pseudo-bicistronic transcripts). While differential expression analysis necessarily requires the use of a p-value cutoff, small changes in transcript stability for a larger number of transcripts may be important. Intriguingly, genes encoding the NMD machinery have been observed to be targets of NMD, suggesting an autoregulatory mechanism for the pathway (Dehecq et al., 2018; Luke et al., 2007; Yepiskoposyan et al., 2011). Similarly, we found both *NMD4* and *EBS1* are NMD-sensitive transcripts, potentially due to a long 3’UTR in the case of the former and uORFs in the case of the latter. It is therefore conceivable that NMD is regulated by external stress or development to evoke functional outcomes (Goetz and Wilkinson, 2017).

Of particular interest, our work establishes a role for NMD in regulating 5’-extended transcripts. The existence of an ensemble of transcript isoforms for a given gene in yeast has been long understood, though often overlooked, in part because the alternative isoforms are often degraded by NMD. Based on our work and that of others, these long transcripts can serve as LUTIs and regulate the expression of downstream genes. Our work focused on the role of LUTIs in nitrogen regulation, and particularly the ensemble of transcripts associated with *DAL5*. We have established that the 5’-extended isoform of *DAL5* is sensitive to NMD and confirmed its function as a LUTI. Our work also suggests the LUTI of *DAL5* not only plays a role in regulating the steady-state levels of the downstream transcript but also the kinetics of the response. This could help the cell more rapidly adjust to changes in sources of nitrogen when it becomes scarce. It also appears to help the cell rapidly shut off production of the Dal5 protein. That 33 AUGs (four of which are expressed) in the LUTI are present to fully suppress protein production from the LUTI (Figure 4E) underscores the likely functional importance for strictly limiting Dal5 protein production under rich nitrogen conditions. It may be that low-level expression of the Dal5 permease is detrimental because it could allow other toxic small molecules into the cell, as suggested by prior work (Cai et al., 2007). Intriguingly, many of the LUTIs we observed in the nitrogen and thiamine regulation pathways, such as *DAL5* (Figures 2D and 5B), *DAL7* (Figures 4C and S5A) *MAK31-PET18* (Figure S4E and 5E), and *THI22* (Figure S5D), are constitutively active. While there are exceptions where the LUTI does turn off under inducing conditions (*DUR3*, data not shown) and *THI6* (Figures S5E), this trend suggests the silencing role for these LUTIs is particularly important.

In principle, the need to degrade these LUTIs and ITIs is not immediately obvious since they do not lead to protein production, and this is supported by our finding that not all LUTIs and ITIs are degraded by NMD. It may be the case that NMD is needed to degrade these transcripts since the uORFs or iORFs that are translated on them encode small peptides that have detrimental outcomes. Alternatively, it may be that the overall mass of these transcripts places an undue burden on the cell by tying up cellular resources. Future work to address these questions is important because PTC mutations are the cause of approximately 11% of inherited genetic disorders (Mort et al., 2008). Many efforts have been reported toward the development of therapeutics to facilitate the readthrough of stop codons, thus inhibiting effects of NMD in the cell (Albers et al., 2023; Coelho et al., 2024; Coller and Ignatova, 2024; Gurzeler et al., 2023; Oltion et al., 2023; Peltz et al., 2013; Sharma et al., 2021). We look forward to additional studies to further unravel the basic role of NMD in the cell and thereby help inform efforts to modulate its effects in human disease.

## Acknowledgements

We thank Alan Hinnebusch, Tom Dever, Jon Lorsch, and Guydosh lab members for useful feedback during the course of this project. We thank Alan Hinnebusch for providing the transcription terminator construct and wig files from published CAGE-Seq data. We thank Elçin Ünal and Gloria Brar for helpful discussions about LUTIs. We thank Hernan Lorenzi for helpful discussions about computational analysis.

This research was supported by the Intramural Research Program of the NIH, the National Institute of Diabetes and Digestive and Kidney Diseases (NIDDK) (DK075132 to N.R.G.).

## Author contributions

D.J.Y. designed experiments, performed high-throughput sequencing and blotting experiments, analyzed the data, and wrote the paper. Y.W. analyzed RNA-Seq data from related yeast species. N.R.G. designed experiments, analyzed data, and wrote the paper.

## SUPPLEMENTAL INFORMATION

### SUPPLEMENTAL FIGURES

**Figure S1.**
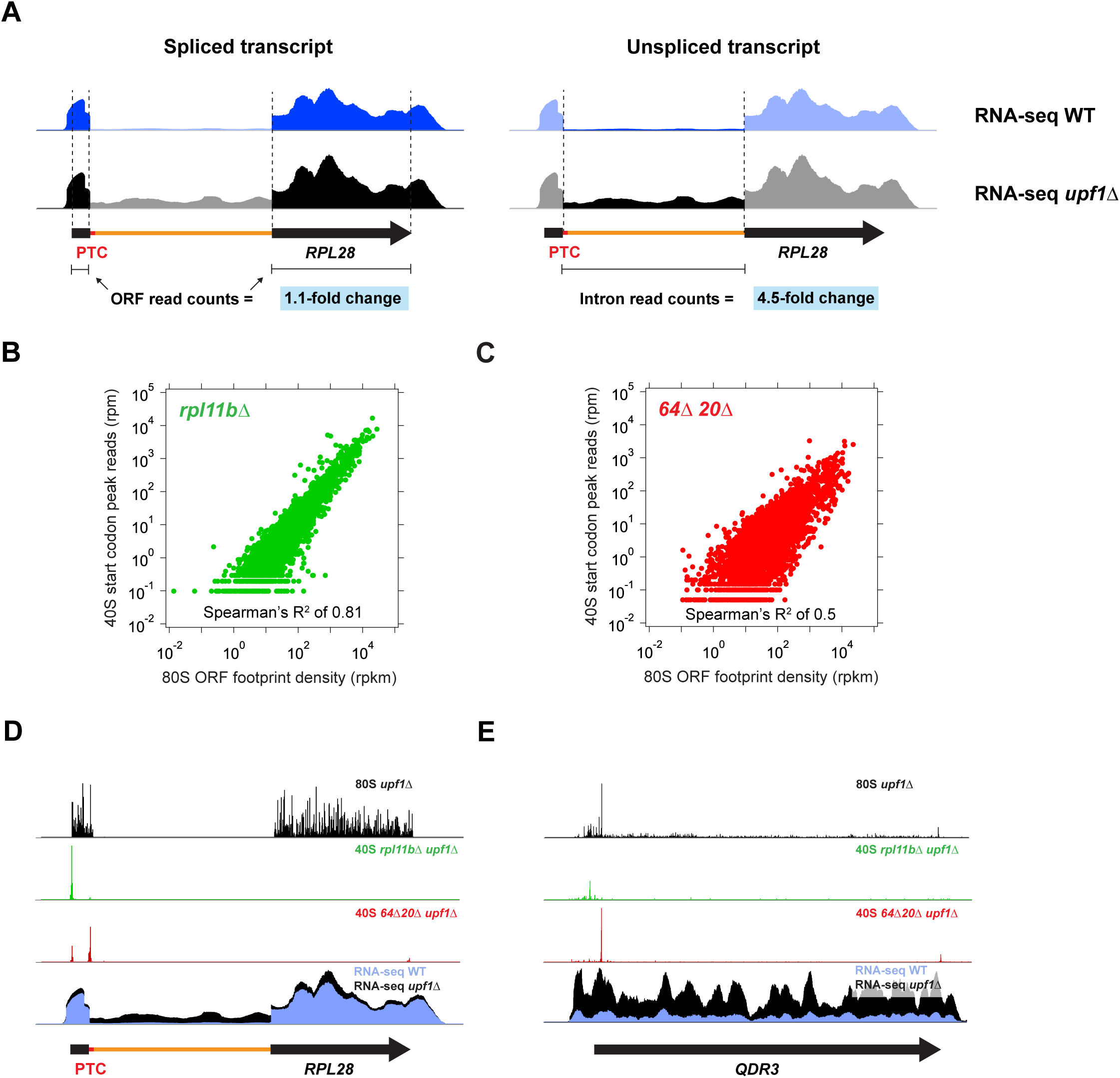
Related to Figures 1 and 2. 40S ribosome profiling reveals translation events on NMD-sensitive transcripts. A) mRNA-Seq data for *RPL28* demonstrating sensitivity to NMD for the intron-containing transcript but not the spliced transcript. B) Correlation of start codon peak height in 40S profiling data from the *rpl11b*Δ strain with 80S ribosome profiling data from the *rpl11b*Δ strain demonstrates utility of using start codon peaks to locate alternative translation events. Spearman R^2^ = 0.81. C) Correlation of stop codon peak height in 40S profiling data from the *tma64*Δ *tma20*Δ strain with 80S ribosome profiling data from the *tma64*Δ *tma20*Δ strain demonstrates utility of using stop codon peaks to locate alternative translation events. Spearman R^2^ = 0.50. D) Sequencing reads for the *RPL28* gene (black arrow corresponds to main ORF exons; red thin line corresponds to the PTC; orange thin arrow corresponds to the intron) show an example of a poorly spliced transcript. 80S ribosome profiling (top) reveals translation into the intron region. 40S profiling (middle tracks) shows peaks at the main ORF start codon and the PTC in the intron and the main stop codon. mRNA-Seq (bottom) shows expression of the intron-containing transcript and its sensitivity to NMD. E) Sequencing reads for the *QDR3* gene in rich media (arrow corresponds to main ORF) showing an example of a short-5’UTR transcript. 80S ribosome profiling (top) reveals translation of an overlapping uORF. 40S profiling (middle tracks) shows peaks at the start and stop codons of the overlapping uORF. mRNA-Seq (bottom) shows expression of the short transcript and its sensitivity to NMD.

**Figure S2.**
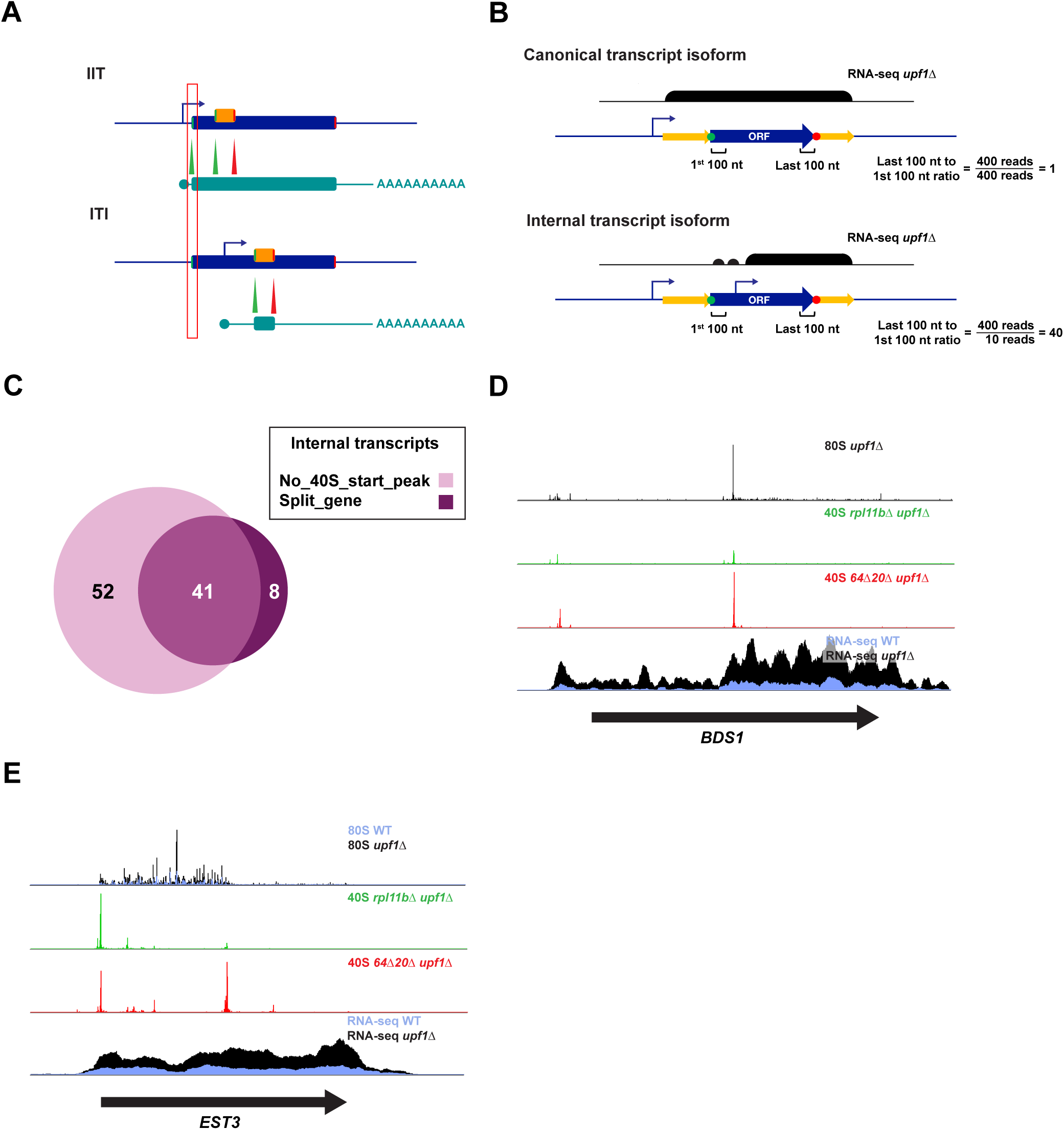
Related to Figure 3. Analysis of translation events within the main ORFs of NMD-sensitive transcripts. A) Differentiating between inefficient initiation transcripts (IITs) and internal transcript isoforms (ITIs). Transcripts that initiate translation inefficiently due to leaky scanning or a short transcript leader exhibit 40S peaks at the main ORF start codon in 40S profiling data whereas ITIs do not (red box). B) Description of the split gene test used to identify ITIs. RNA-Seq reads from the first and last 100 nt of the main ORF are compared. Those genes with >2-fold more reads in the last 100 nt are considered ITIs. C) Comparison of genes identified as ITIs using the absence of a 40S profiling peak at the main ORF start codon vs the split-gene test. Results from both tests were combined given the substantial overlap. D) Sequencing reads for the *BDS1* gene in rich media (arrow corresponds to main ORF) showing an example of a case where both a 5’-extended transcript and ITI are expressed. 80S ribosome profiling (top) shows uORF and iORF translation. 40S profiling (middle tracks) shows locations of start and stop codons on the uORF and iORF. mRNA-Seq (bottom) shows expression of the 5’-extended transcript and ITI and their sensitivity to NMD. E) Sequencing reads for the *EST3* gene in rich media (arrow corresponds to main ORF) showing an example of a case where a programmed frameshift leads to NMD. 80S ribosome profiling (top) shows translation up to the site of frameshift. Low frameshift efficiency results in most 80S footprints terminating just after the site of the frameshift. 40S profiling (middle tracks) shows the location of the stop codon after the frameshift site. mRNA-Seq (bottom) shows expression of the transcript and its sensitivity to NMD.

**Figure S3.**
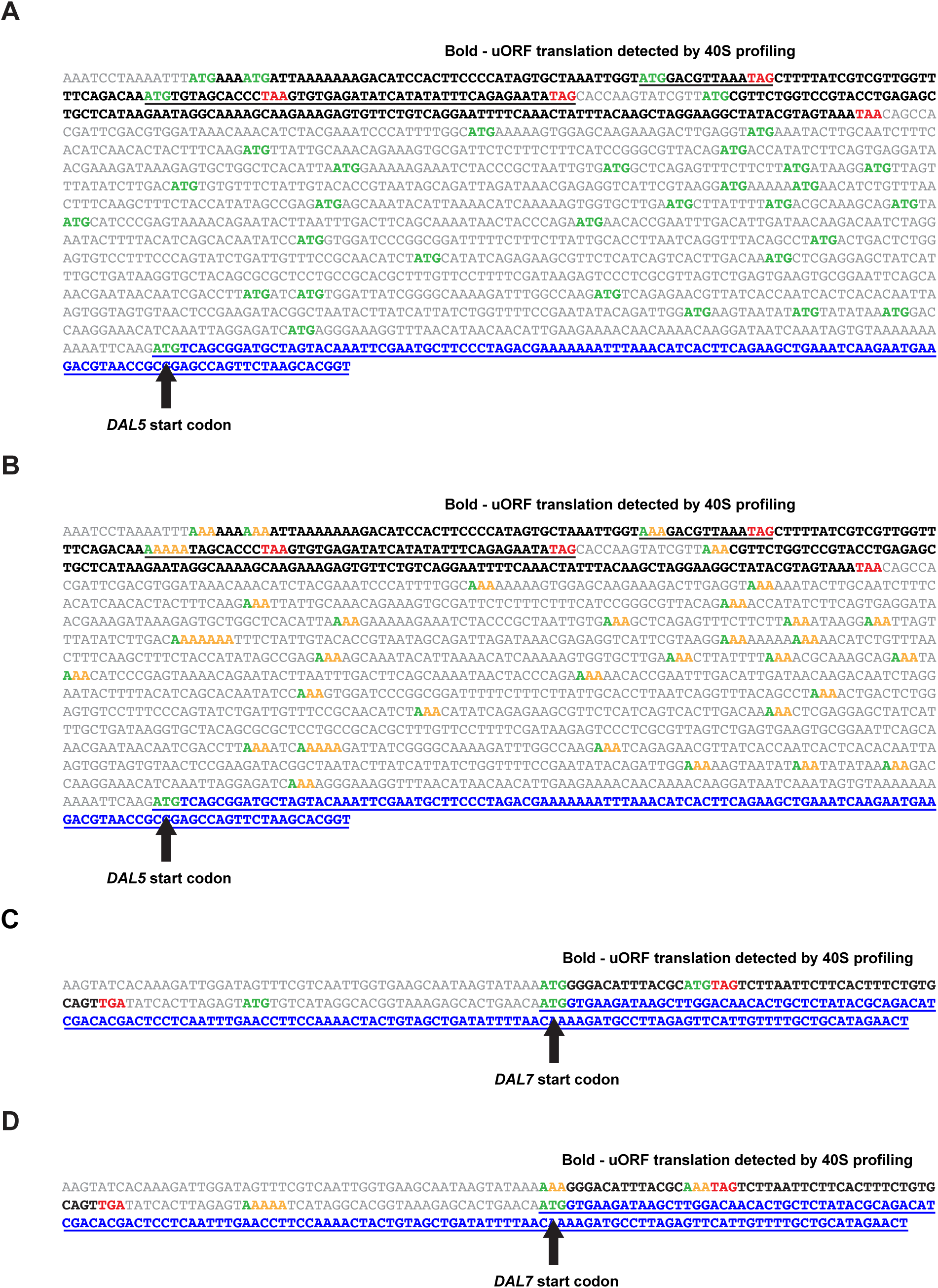
Related to Figure 4. Map of uORFs on the *DAL5* and *DAL7* transcripts. A) Location of the 33 AUG codons in the 5’-extended transcript of *DAL5*. Translated uORFs (detected by 40S ribosome profiling) are shown in bold. B) Location of the 33 AUG codons in the 5’-extended transcript of *DAL5* shown mutated to AAA for northern- and western-blotting experiments. Translated uORFs (detected by 40S ribosome profiling) shown in bold. C) Same as A for *DAL7*. D) Same as B for *DAL7*. Underlining is used throughout to help distinguish overlapping ORFs.

**Figure S4.**
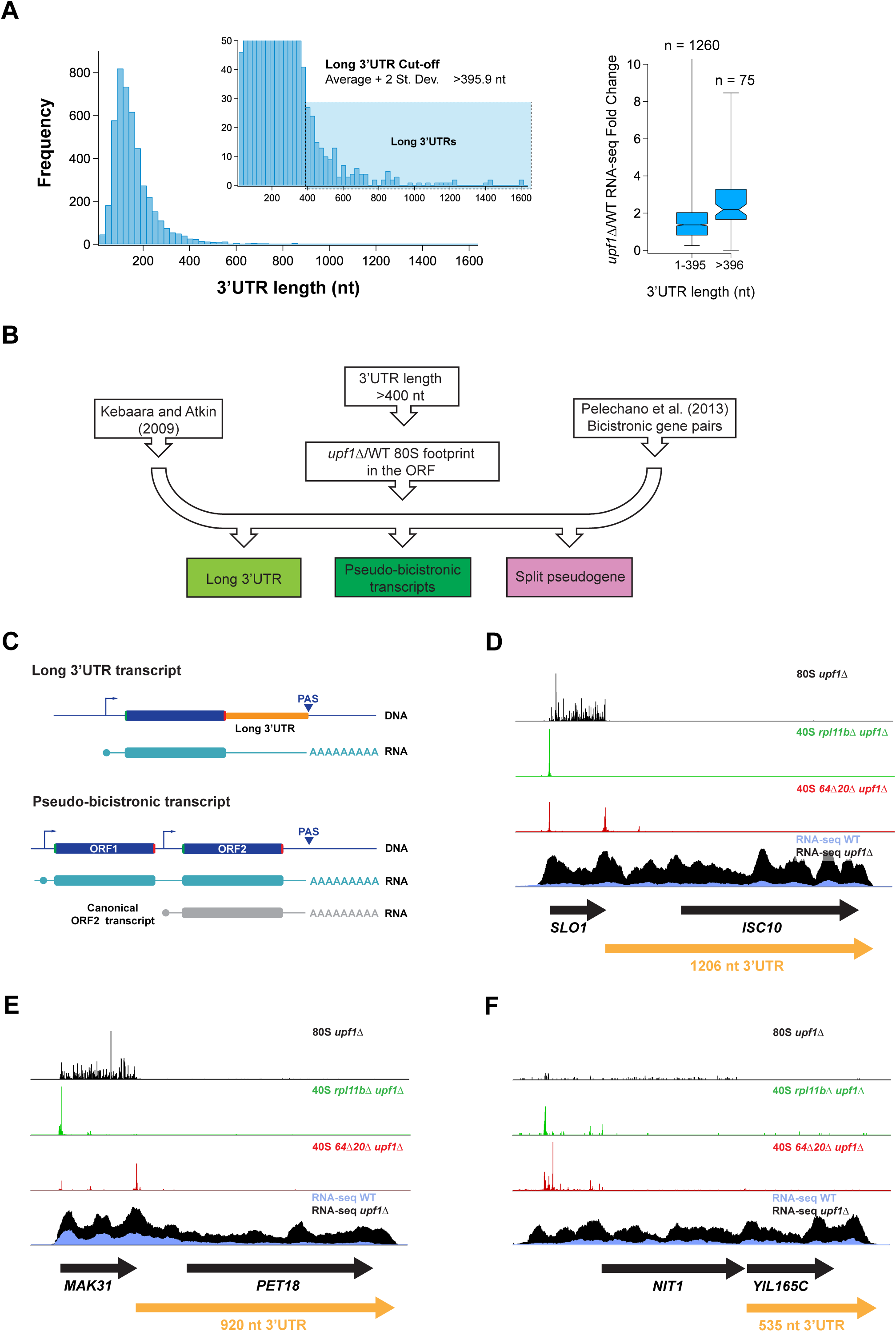
Related to Figure 1. Analysis and assessment of genes with long 3’UTRs associated with NMD-sensitive transcripts. A) (Left) Distribution of annotated most abundant 3’UTR lengths in yeast grown in glucose. (Right) Analysis of NMD sensitivity for transcripts with long (>396 nt) 3’UTRs vs those with short annotated 3’UTRs. On average, long 3’UTRs are more sensitive to NMD. NMD-sensitive genes included with FDR<0.01. B) Pipeline used to determine likely cases where long 3’UTRs may sensitize transcripts to NMD and assign according to 3 output categories. Apart from the 3 inputs of gene names, the 80S footprint requirement confirms that the main ORF of the transcript targeted to NMD is translated. C) Potential architecture of yeast genes with long 3’UTRs: a long 3’UTR is encoded for some genes (top) or a pseudo-bicistronic transcript results when the transcription terminator at the end of ORF1 is either deleted or poor, leading to (at least) some production of a transcript covering both genes that acts like a transcript with long 3’UTR (bottom). D) Sequencing reads for the *SLO1-ISC10* pseudo-bicistronic gene in rich media (black arrow corresponds to main ORFs) show this transcript is a candidate long 3’UTR transcript. 80S ribosome profiling (top) reveals translation of the first main ORF. 40S profiling (middle tracks) shows locations of start and stop codons of the first main ORF. mRNA-Seq (bottom) shows expression of pseudo-bicistronic transcript and its sensitivity to NMD. Yellow arrow indicates effective 3’UTR of *SLO1* transcript. E) Sequencing reads for the *MAK31-PET18* pseudo-bicistronic gene in rich media (black arrow corresponds to main ORFs) show this transcript is a candidate long 3’UTR transcript. 80S ribosome profiling (top) reveals translation of the first main ORF. 40S profiling (middle tracks) shows locations of start and stop codons of the first main ORF. mRNA-Seq (bottom) shows expression of pseudo-bicistronic transcript and its sensitivity to NMD, as well as some abundance of a transcript that terminates between the two ORFs. Yellow arrow indicates effective 3’UTR of *MAK31* transcript. F) Sequencing reads for the *NIT1-YIL165C* pseudogene transcript in rich media (black arrow corresponds to main ORFs) show this transcript is a candidate long 3’UTR transcript. 80S ribosome profiling (top) shows some translation of the first main ORF. 40S profiling (middle tracks) shows locations of start and stop codons on first main ORF. mRNA-Seq (bottom) shows expression of psuedogene transcript and its sensitivity to NMD. We note that this transcript also exhibits uORFs on an apparent 5’-extended transcript (see Figure S6A). Yellow arrow indicates effective 3’UTR of *NIT1* transcript.

**Figure S5.**
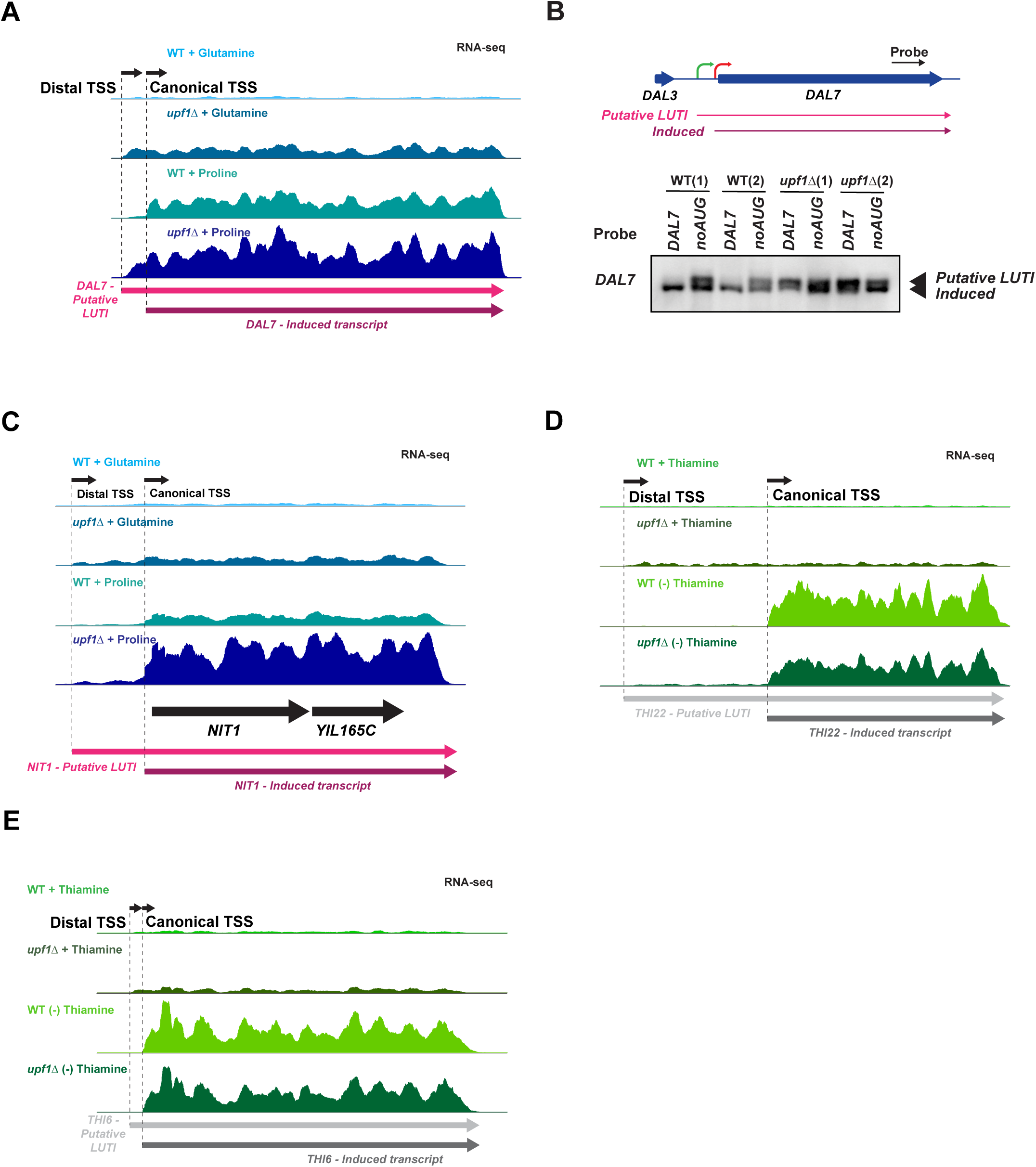
Related to Figure 5. Characterization of regulated transcript pairs that include a long NMD-sensitive (putative LUTI) transcript and a downstream (canonical) transcript. A) mRNA-Seq reads mapped to the *DAL7* gene in order from top to bottom: WT in nitrogen-rich media, *upf1*Δ in nitrogen-rich media, WT in nitrogen-poor media, *upf1*Δ in nitrogen-poor media. Black arrow on top show approximate TSSs. Colored arrows at bottom show transcripts. Data reveal that the NMD-sensitive, putative LUTI is present under both media conditions and the existence of a shorter canonical NMD-insensitive transcript is induced under poor nitrogen. B) Northern analysis confirms expression for *DAL7* of both a putative LUTI transcript and a short canonical induced transcript under poor nitrogen conditions. The 5’ uORFs confer NMD sensitivity to only the long transcript. C) mRNA-Seq reads mapped to the *THI22* gene in order from top to bottom: WT in media with thiamine, *upf1*Δ in media with thiamine, WT in media without thiamine, *upf1*Δ in media without thiamine. Black arrow on top show approximate TSSs. Arrows at bottom show transcripts. Data reveal that the NMD-sensitive putative LUTI is present under both media conditions and the existence of a shorter canonical NMD-insensitive transcript that is induced in the absence of thiamine. D) mRNA-Seq reads mapped to the *THI6* gene in order from top to bottom: WT in media with thiamine, *upf1*Δ in media with thiamine, WT in media without thiamine, *upf1*Δ in media without thiamine. Black arrow on top show approximate TSSs. Arrows at bottom show transcripts. Data reveal that the NMD-sensitive putative LUTI is present only in the presence of thiamine and the NMD-insensitive induced transcript is present only in the absence of thiamine. E) mRNA-Seq reads mapped to the *NIT1*/*YIL164C-YIL165C* pseudogene in order from top to bottom: WT in nitrogen-rich media, *upf1*Δ in nitrogen-rich media, WT in nitrogen-poor media, *upf1*Δ in nitrogen-poor media. Black arrow on top show approximate TSSs. Colored arrows at bottom show transcripts. Data reveal an NMD-sensitive putative LUTI is present under both media conditions and an NMD-sensitive pseudogene transcript containing a PTC is induced under poor nitrogen.

**Figure S6.**
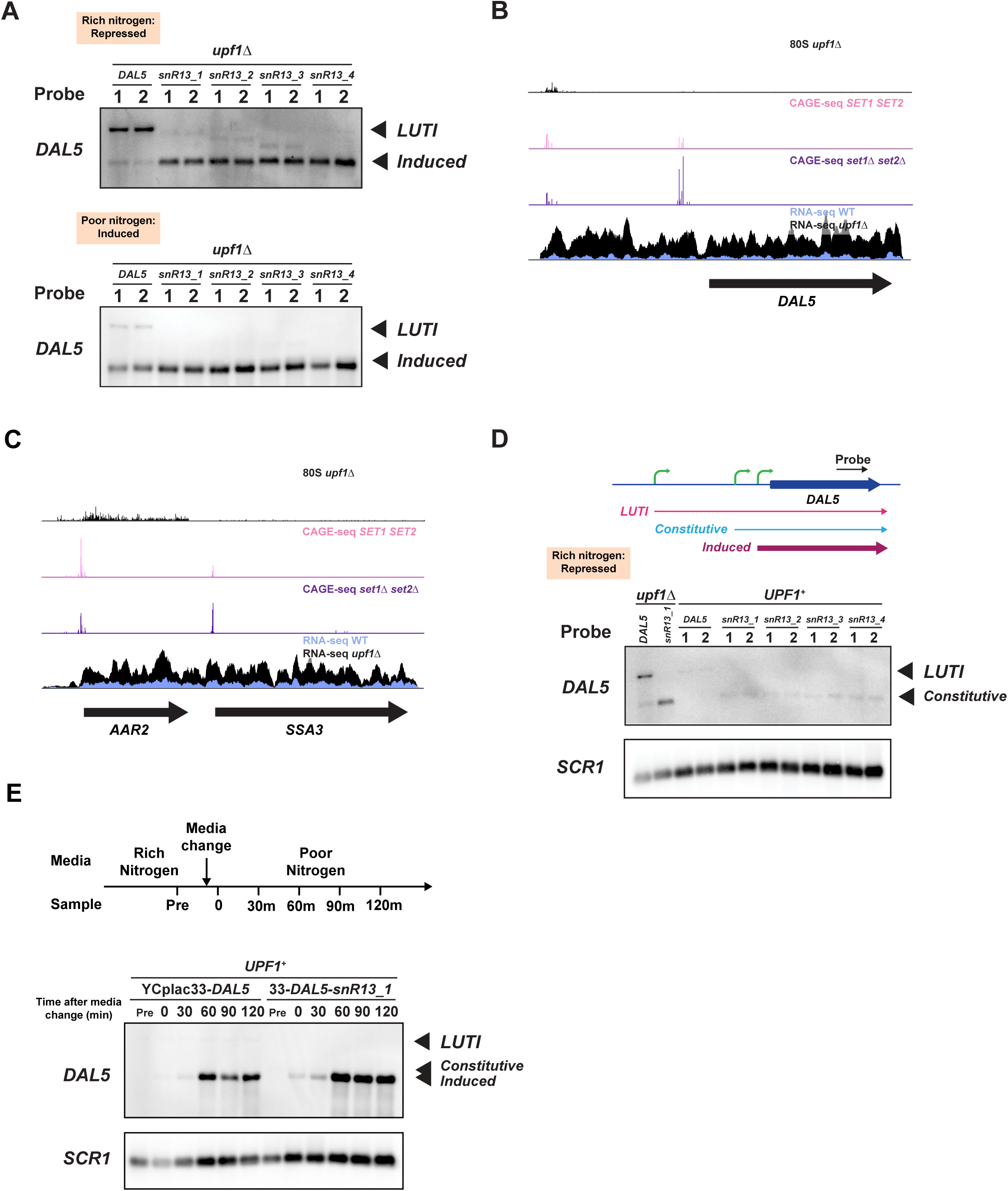
Related to Figure 6. The *DAL5* LUTI represses the constitutive transcript under rich nitrogen and the induced transcript under poor nitrogen. A) Northern analysis of *DAL5* shows premature termination of LUTI transcript derepresses downstream transcripts under both rich and poor nitrogen conditions. B) CAGE-Seq and mRNA-Seq reads mapped to the *DAL5* gene in order from top to bottom: *upf1*Δ 80S ribosome profiling (top) in nitrogen-rich media, WT (*SET1 SET2*) CAGE-Seq in nitrogen-rich media, *set1*Δ *set2*Δ CAGE-Seq in nitrogen-rich media, WT mRNA-Seq in nitrogen-rich media, and *upf1*Δ mRNA-Seq in nitrogen-rich media. Arrow corresponds to the main ORF. CAGE-Seq data reveal locations of 5’-extended (LUTI) TSS and the TSS of the constitutive transcript that is depressed in the absence of *SET1* and *SET2*. C) CAGE-Seq and mRNA-Seq reads mapped to the *AAR2-SSA3* pseudo-bicistronic gene in order from top to bottom: *upf1*Δ 80S ribosome profiling (top) in nitrogen-rich media, WT (*SET1 SET2*) CAGE-Seq in nitrogen-rich media, *set1*Δ *set2*Δ CAGE-Seq in nitrogen-rich media, WT mRNA-Seq in nitrogen-rich media, and *upf1*Δ mRNA-Seq in nitrogen-rich media. Arrow corresponds to the main ORF. CAGE-Seq data reveals the locations of the *AAR2* TSS and the TSS of the *SSA3* transcript that is depressed in the absence of *SET1* and *SET2*. D) Northern analysis of *DAL5* shows that under WT (*UPF1*^+^) conditions the constitutive transcript that is derepressed upon premature termination of the LUTI transcript is NMD-sensitive. E) Repeat of derepression experiment in 7A performed for longer timescale to establish endpoint. In blots above, 2 replicates are shown for each condition.

**Tables S1, S2, S6, and S9 are available as downloadable files. Tables S3-S5 and S7-S8 are below.**

**Table S3:** Plasmids used in this study, related to STAR Methods.

| **Plasmid** | **Description** | **Source** |
| --- | --- | --- |
| pAG32 | pFA6-natMX4; *NAT* deletion cassette | Goldstein and McCusker, 1999 |
| pFA6-kanMX4 | *KAN* deletion cassette | Wach et al. 1994 |
| YCplac33 | sc<sup>1</sup> *URA3* | Gietz and Sugino 1988 |
| pDY243 | YCplac33-DAL5 | This study |
| pDY236 | YCplac33-DAL7 | This study |
| pDY270 | pUC57-DAL5_LUTI-noAUG | Genscript USA, Inc. |
| pDY284 | YCplac33-DAL5_LUTI-noAUG | This study |
| pDY286 | YCplac33-DAL7_LUTI-noAUG | This study |
| pDY290 | YCplac33-DAL5-snR13_1 | This study |
| pDY292 | YCplac33-DAL5-snR13_2 | This study |
| pDY294 | YCplac33-DAL5-snR13_3 | This study |
| pDY296 | YCplac33-DAL5-snR13_4 | This study |

<sup>1</sup>sc, single-copy

**Table S4:**
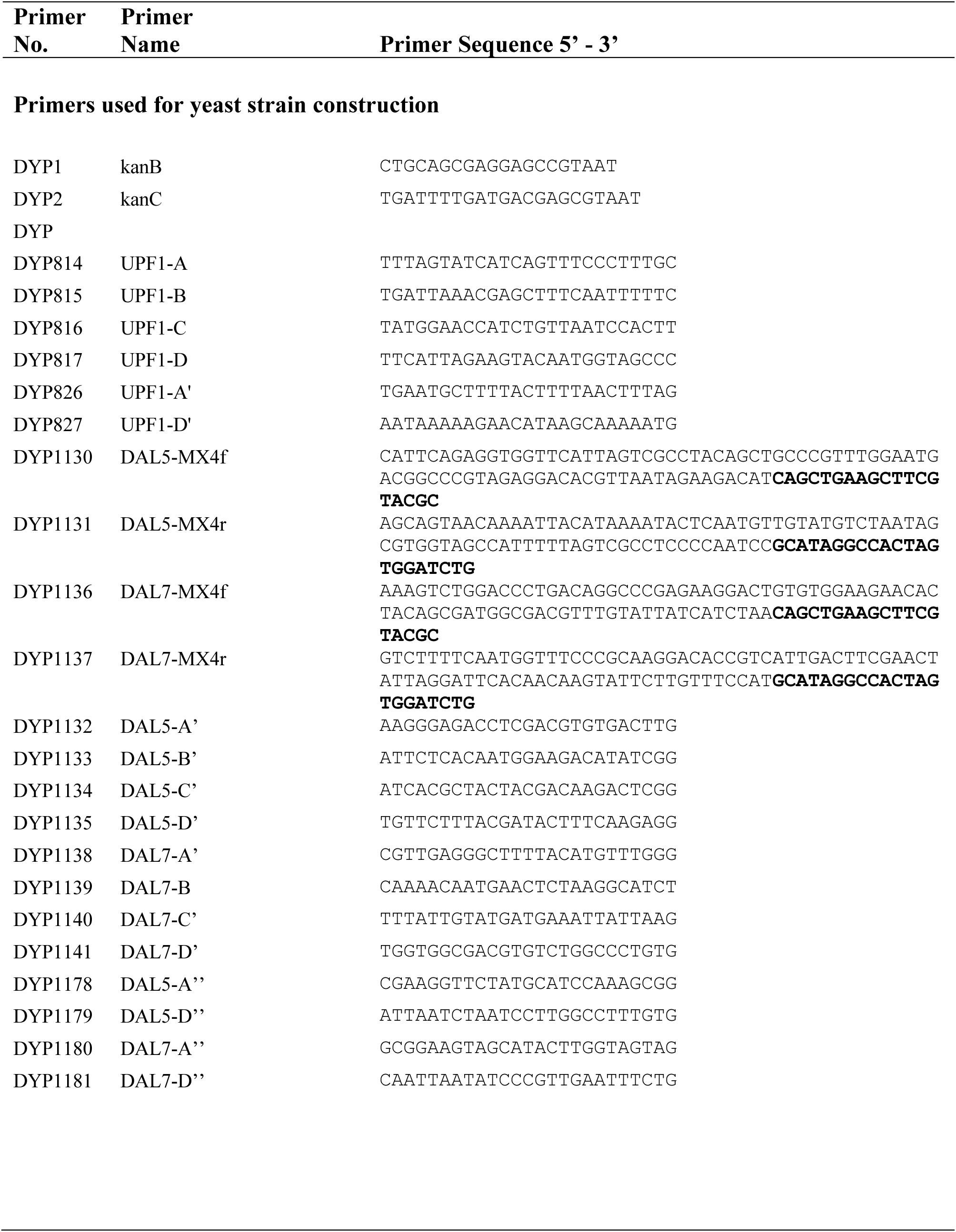
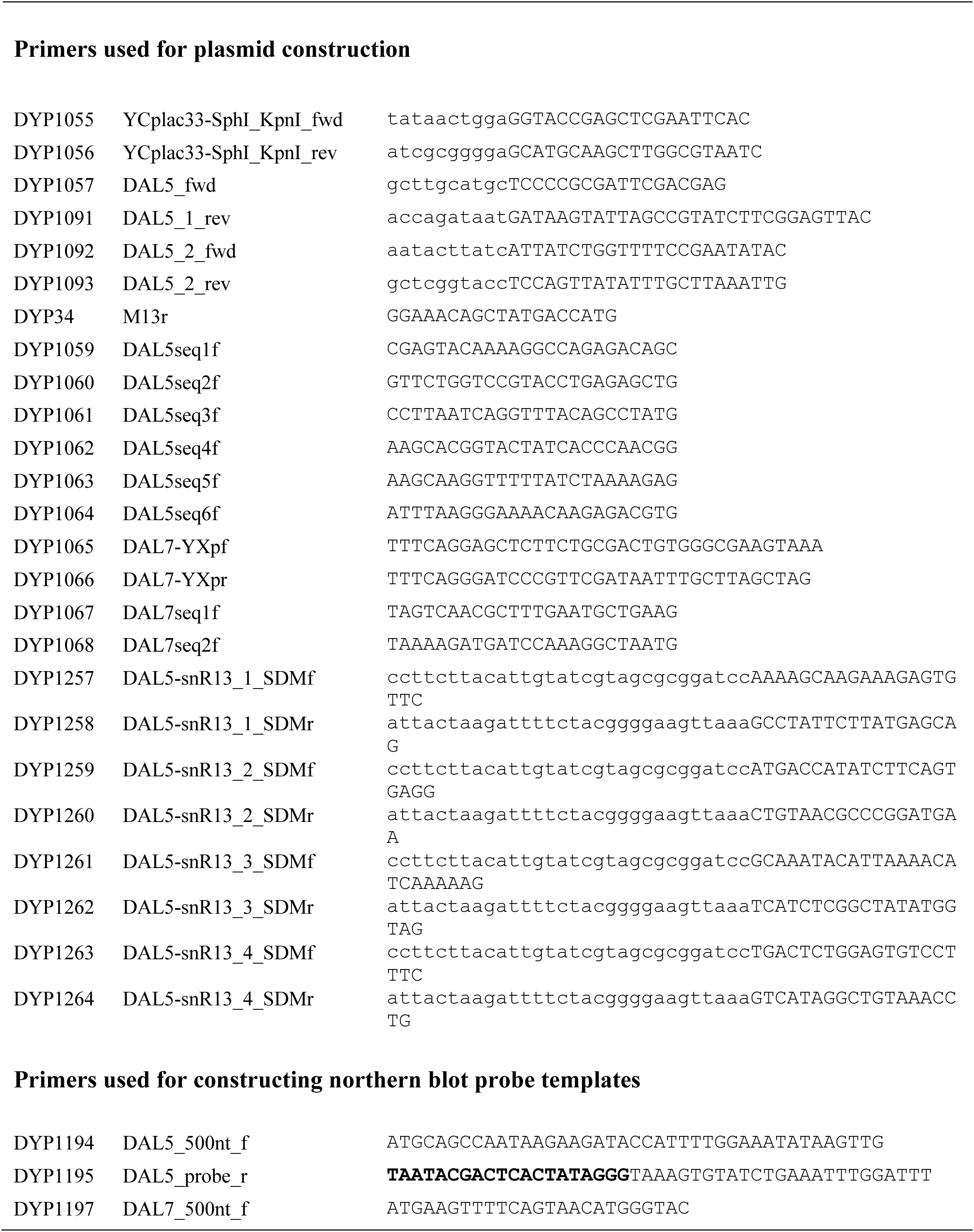
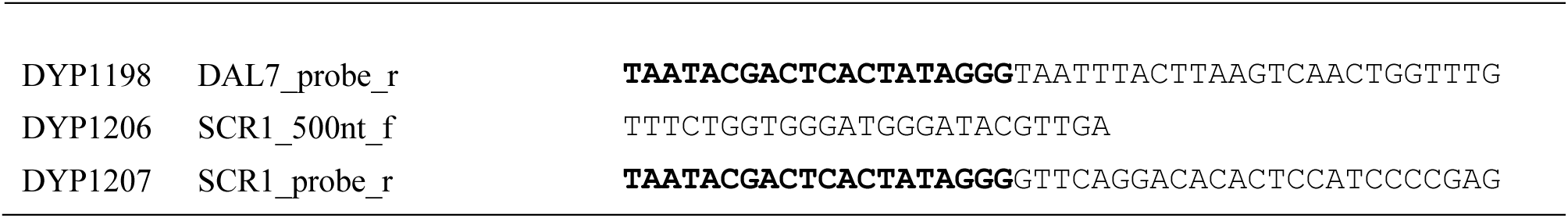
Oligonucleotides used in this study, related to STAR Methods.

**Table S5:**
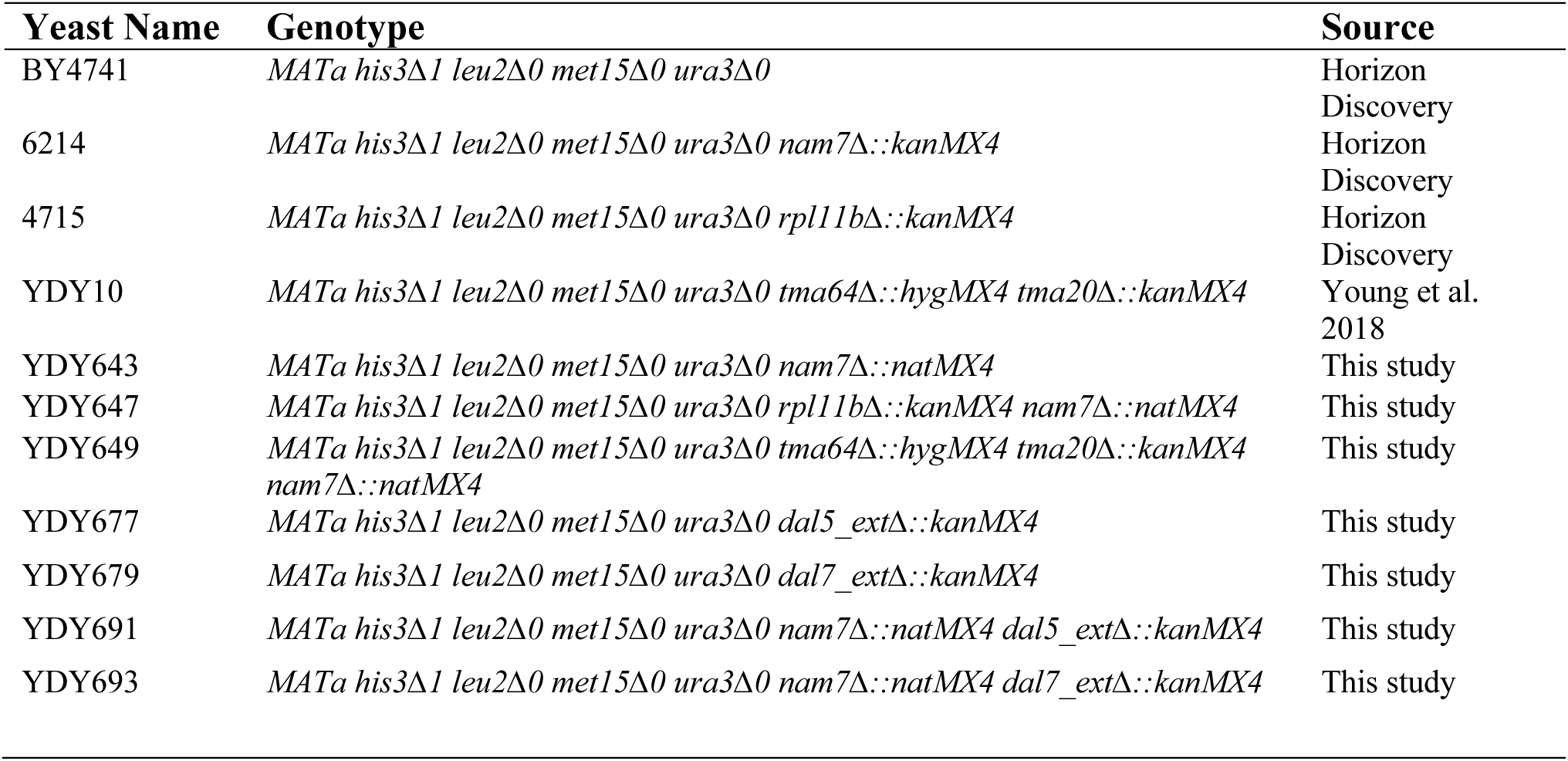
Yeast Strains used in this study, related to Methods.

**Table S7:** 40S ribosome profiling datasets from previous papers, related to Figure S1 and Methods.

| **Sample name** | **Description** | **Reference** | **GEO Number** | **Number mapped reads** |
| --- | --- | --- | --- | --- |
| DY135Ffx1<br>83Ffx | *rpl11b*Δ (2 rep's pooled) | Young et al. (2021) | GSM4339086 | 6,568,093 |
| DY136Ffx1<br>84Ffx | *tma64*Δ/*tma20*Δ (2 rep's pooled) | Young et al. (2021) | GSM4339087 | 12,482,481 |

**Table S8:**
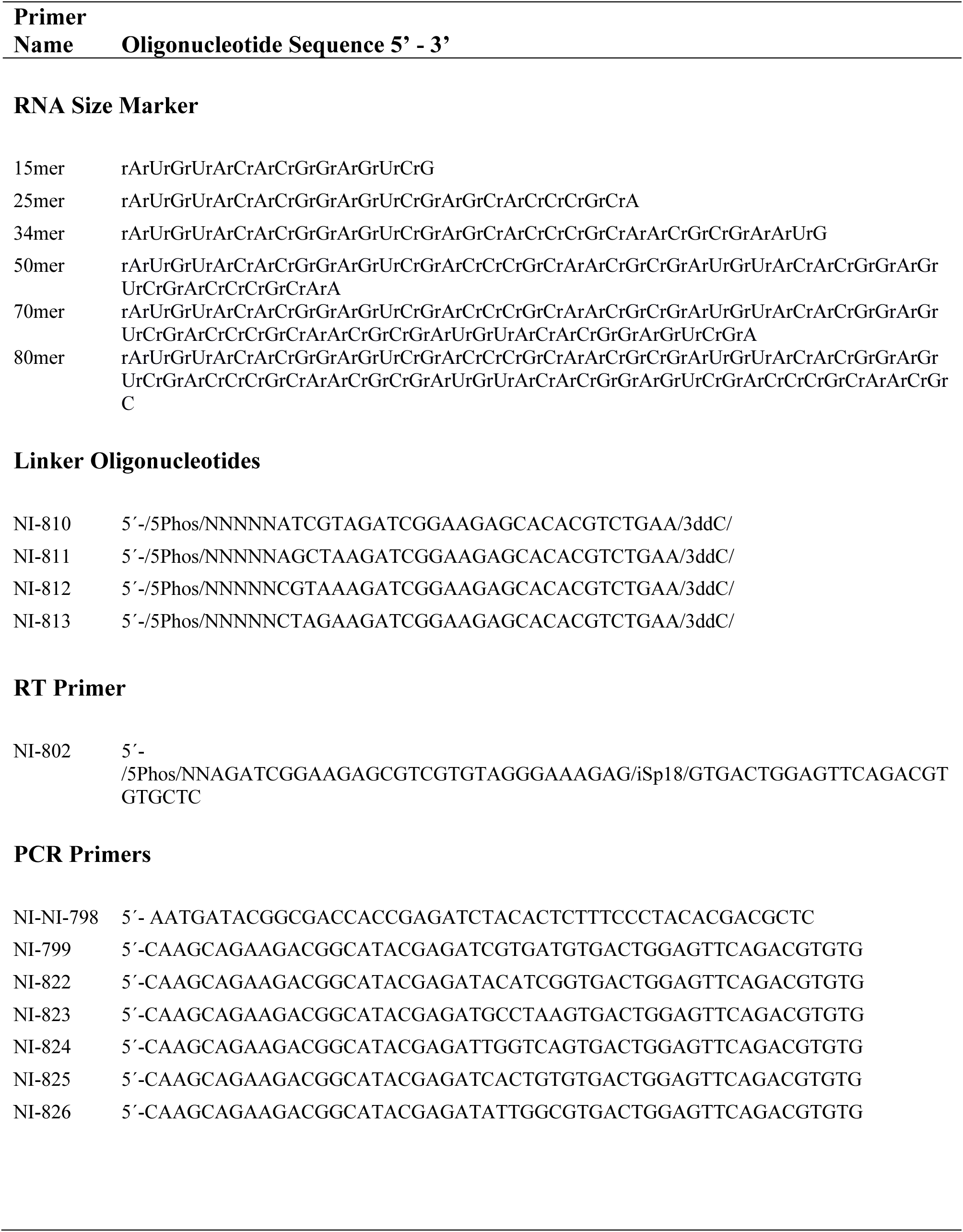
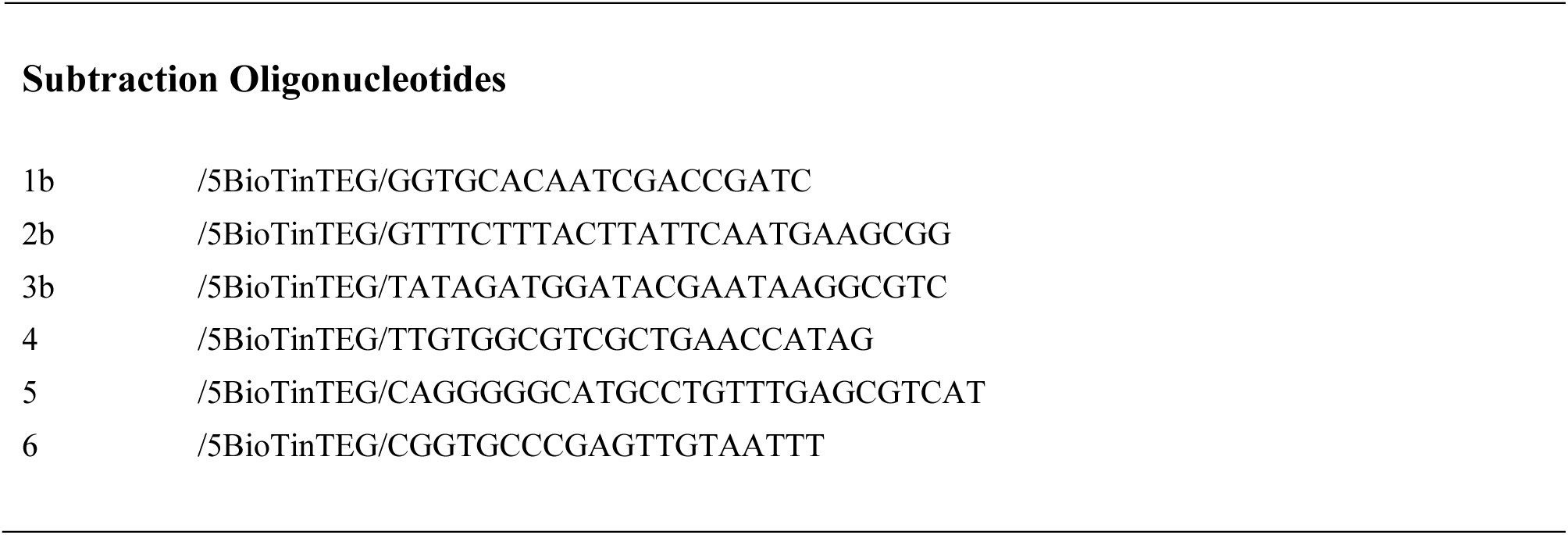
Oligonucleotides used for ribosome profiling, related to Methods.

## METHODS

### Plasmid Constructions

Plasmids used in this study are listed in Supplemental Table S3. The primers used for plasmid construction are listed in Supplemental Table S4.

To manipulate the *DAL5* 5’-extended transcript a 5 kb region of chromosome X (chrX:717,000-721,999) encompassing the *DAL5* ORF, and the entirety of the 5’extended transcript with a 1kb upstream margin to ensure capture of the uncharacterized 5’-extended transcript promoter was cloned into YCplac33 (sc *URA3* plasmid).

Plasmid pDY243 (YCplac33-*DAL5*) was constructed using the NEBuilder HiFi DNA Cloning Kit (NEB; E5520S). Briefly, the YCplac33 backbone was PCR amplified from YCplac33 using primers YCplac33-SphI_KpnI_fwd and YCplac33-SphI_KpnI_rev. The *DAL5* extended 5 kb region was PCR amplified in two 2.5 kb sections from yeast gDNA using primer pairs DAL5_fwd/DAL5_1_rev and DAL5_2_fwd/DAL5_2_rev. The three PCR products were purified and assembled according to the manufacturer’s instructions. Correct integration of the *DAL5* extended 5 kb region into the YCplac33 backbone was confirmed by DNA sequencing.

To manipulate the *DAL7* long transcript a 2.259 kb region of chromosome IX (chrIX:412771-415029) encompassing the *DAL7* ORF, the NMD-sensitive uORF-containing transcript, and the *DAL7* promoter was cloned into YCplac33 (sc *URA3* plasmid).

Plasmid pDY236 (YCplac33-*DAL7*) was constructed by PCR amplifying the *DAL7* gene from yeast genomic DNA using DAL7-YXpf and DAL7-YXpr, digesting with SacI-HF and BamHI-HF and inserting the resulting restriction fragment between the SacI and BamHI sites of YCplac33. Correct insertion of the *DAL7* extended region into the YCplac33 backbone was confirmed by DNA sequencing.

To construct plasmids pDY284 and pDY286 (YCplac33-*DAL5_LUTI-noAUG* and YCplac33-*DAL7_LUTI-noAUG*) fragments of YCplac33-DAL5 and YCplac33-DAL7 bounded by restriction sites unique to each plasmid were synthesized where all the AUG codons present in the 5’UTR of each LUTI were mutated to AAA (Supplemental Document - LUTI replacement windows). The 2840 bp *DAL5_LUTI-noAUG* fragment was synthesized by GenScript USA, Inc. and cloned into pUC57 (pUC57-*DAL5_LUTI-noAUG*). The 1180 bp *DAL7_LUTI-noAUG* fragment was synthesized as a gBlock by Integrated DNA Technologies, Inc. pUC57-*DAL5_LUTI-noAUG* was digested with SphI-HF and SacII and the resulting 2765 bp restriction fragment was inserted between the SphI and SacII sites of YCplac33-DAL5 to produce plasmid pDY284. The *DAL7_LUTI-noAUG* gBlock was digested with EcoRI-HF and KpnI-HF and the resulting 1115 bp restriction fragment was inserted between the EcoRI and KpnI sites of YCplac33-DAL7 to produce plasmid pDY286. Correct insertion of the fragments was confirmed by DNA sequencing.

Plasmids pDY290 (YCplac33-*DAL5-snR13_1*), pDY292 (YCplac33-*DAL5-snR13_2*), pDY294 (YCplac33-*DAL5-snR13_3*), and pDY296 (YCplac33-*DAL5-snR13_4*) were constructed using the Q5 Site-directed mutagenesis kit (NEB; E0554S) and primer pairs DAL5-snR13_1_SDMf / DAL5-snR13_1_SDMr, DAL5-snR13_2_SDMf / DAL5-snR13_2_SDMr, DAL5-snR13_3_SDMf / DAL5-snR13_3_SDMr, and DAL5-snR13_4_SDMf / DAL5-snR13_4_SDMr respectively.

### Yeast Strain Construction

All *Saccharomyces cerevisiae* strains used in this study are derived from the BY4741 background. They were maintained on either YPD plates or SC-Ura plates for transformants. Yeast strains used in this study are listed in Table S5. The primers used for strain construction and verification are listed in Table S4.

The *rpl11b1*Δ *nam7*Δ (*upf1*Δ) and *tma64*Δ *tma20*Δ *nam7*Δ (*upf1*Δ) double and triple deletion strains (YDY647 and YDY649) were constructed in two steps. First, the *nam7*Δ*::kanMX4* deletion allele in strain 6214 was converted to *nam7*Δ*::natMX4*, by transformation with BamHI/SpeI digested pAG25 (carrying *natMX4*), followed by selection on YPD containing 100 μg/mL ClonNat to produce strain YDY643. The *nam7*Δ*::natMX4* allele was then PCR-amplified from strain YDY643 and transformed into strains 4715 (*rpl11b*Δ) and YDY10 (*tma64*Δ *tma20*Δ) to produce strains YDY647 and YDY649. PCR amplification from chromosomal DNA was used to confirm correct integrations.

WT and *upf1*Δ strains were constructed where the chromosomal regions cloned into YCplac33-*DAL5* and YCplac33-*DAL7* were deleted to avoid signal from endogenous transcripts.

The *dal5_ext*Δ and *dal7_ext*Δ strains (YDY677 and YDY679) were constructed by PCR amplifying the *kanMX4* deletion cassette from pFA6-*kanMX4* using the DAL5-MX4f/DAL5-MX4r and DAL7-MX4f/DAL7-MX4r primer pairs. The purified PCR products were transformed into BY4741 and selected on YPD containing 200 μg/mL G418 to produce strains YDY677 and YDY679. PCR amplification from chromosomal DNA was used to confirm correct integrations.

The *dal5_ext*Δ *upf1*Δ and *dal7_ext*Δ *upf1*Δ strains (YDY691 and YDY693) were constructed by PCR amplifying the *dal5_ext*Δ*::kanMX4* and *dal7_ext*Δ*::kanMX4* alleles from YDY677 and YDY679. The purified PCR products were transformed into YDY643 (*nam7*Δ::*natMX4*) and selected on YPD containing 200 μg/mL G418 to produce strains YDY691 and YDY693. PCR amplification from chromosomal DNA was used to confirm correct integrations.

### RNA-seq and Ribosome profiling

The RNA-seq and ribosome profiling datasets generated for this paper, including number of reads and alignment statistics, are described in >Supplemental Table S6. Previously published 40S ribosome profiling datasets used in this paper are listed in Supplemental Table S7.

### RNA-seq

#### Growth of yeast for RNA-seq

For the initial RNA-seq studies to determine changes in mRNA levels in the *upf1*Δ deletion strain, BY4741 (WT) and 6214 (*nam7*Δ [*upf1*Δ]) were grown in YPD. For growth in rich nitrogen (glutamine) and poor nitrogen (proline) conditions, BY4741 (WT) and 6214 (*nam7*Δ [*upf1*Δ]) were grown in YNB (without amino acids or ammonia) with either glutamine or proline (0.1% final concentration), glucose (3%) and the appropriate supplements to cover auxotrophic requirements (300 μM histidine-HCl, 2 mM Leucine, 1 mM Methionine, and 200 μM Uracil). For growth with and without thiamine, BY4741 (WT) and 6214 (*nam7*Δ [*upf1*Δ]) were grown in YNB (without amino acids or ammonia or thiamine) (Formedium Ltd; CYN4801), ammonium sulfate (0.5%), glucose (2%) and the appropriate supplements to cover auxotrophic requirements (300 μM histidine-HCl, 2 mM Leucine, 1 mM Methionine, and 200 μM Uracil). Thiamine (0.4 mg/L) was added to create the media with thiamine. To induce the UPR, BY4741 (WT) and 6214 (*nam7*Δ [*upf1*Δ]) were grown in YPD to an OD_600_ of 0.4 and treated with 5 mM DTT for 1 h. Cells were grown in 750 mL of media in a 2 L flask to a final OD_600_ of 0.6, fast filtered, and frozen in liquid nitrogen.

#### Preparation of RNA-seq libraries

Cell scrapes were resuspended in lysis buffer (8.4 mM EDTA, 60 mM Sodium acetate) and total RNA was purified using hot phenol chloroform. 5 μg of RNA was fragmented in 2x Fragmentation solution (12 mM Na_2_CO_3_, 88 mM NaHCO_3_, 2mM EDTA, pH 9.2) at 95 °C for 35 min and purified using an oligo clean & concentrator kit (Zymo Research; D4060). 50-70 nt fragments were size selected (see Table S8 for size standards) from a 15% TBE-Urea gel.

Construction of sequencing libraries was performed using a protocol based on the ribosome profiling method described in (McGlincy and Ingolia, 2017). The purified RNA fragments were dephosphorylated using PNK (NEB; M0201L) and ligated to pre-adenylated linkers (Table S6) containing a randomized 5 nt Unique Molecular Index (UMI) and a 5 nt sample barcode unique for each sample using truncated T4 RNA ligase 2 (K227Q) (NEB; M0351L). The linkers were pre-adenylated using a 5’ DNA adenylation kit (NEB; E2610L). Unligated linker was removed from the ligation reaction by addition of 5’ deadenylase (NEB; M0331S) and RecJ exonuclease (Biosearch Technologies; RJ411250). Ligated samples were pooled and purified using an oligo clean & concentrator kit (Zymo Research; D4060). Ribosomal RNAs were removed from the pooled samples using the QIAseq FastSelect – rRNA Yeast Kit (Qiagen; 334215). The pooled samples were reverse transcribed using the RT primer NI-802 (Table S8) containing a randomized 2 nt UMI, and Superscript III (Invitrogen; 18080044). The reverse transcribed footprints were separated from unutilized RT primer on a 10% TBE-Urea gel and circularized using CircLigase ssDNA Ligase (Biosearch Technologies; CL4115K). The circularized libraries were amplified by PCR using Phusion DNA Polymerase (ThermoFisher Scientific; F530L) to add unique 6 nt (“Illumina”) barcodes for index sequencing and common Illumina primer and flow cell binding regions. Library quality and concentration was assessed by BioAnalyzer using the High Sensitivity DNA Kit (Agilent; 5067-4626). Libraries were pooled, and sequencing was performed on an Illumina machine at the NHLBI DNA Sequencing and Genomics Core at NIH (Bethesda, MD).

### 40S ribosome profiling

40S ribosome profiling was performed as described in Young et al. (2021).

#### Growth of yeast for 40S ribosome profiling

YDY647 (*rpl11b1*Δ *nam7*Δ [*upf1*Δ]) and YDY649 (*tma64*Δ *tma20*Δ *nam7*Δ [*upf1*Δ]) were grown in YPD. Cells were grown in 1200 mL of media in a 6 L flask to a final OD_600_ of 1.5.

#### Formaldehyde cross-linking

For each culture, 600 mL of cells was poured into two 1 L precooled centrifuge bottles containing 150g of ice and 16.6 mL of 37% formaldehyde (Sigma; 252549). The centrifuge bottles were mixed by inversion and placed on ice for 1 h, with mixing every 15 min. Cross-linking was stopped by the addition of 30 mL of 2.5 M Glycine (Sigma; G7126). The cells were centrifuged at 4,000 rpm for 20 min at 4 °C, resuspended in 10 mL of water, and combined into one 50 mL conical tube. The cells were centrifuged at 4,000 rpm for 5 min at 4 °C and resuspended in 2 mL of lysis buffer (20 mM Tris pH8, 140 mM KCl, 1.5 mM MgCl_2_, 1% Triton X-100). The resuspended cells were beaded into liquid nitrogen, transferred to a pre-chilled 50 mL conical tube, and stored in a -80 °C freezer.

#### Preparation of 40S footprint libraries

Cells were lysed in a Retsch Cryomill (Retsch 20.749.0001). The milled cells were transferred to a 50 mL conical tube, thawed, and spun at 3000g for 5 min at 4 °C. The supernatant was transferred to 1.5 mL Eppendorf tubes and spun at full speed for 10 min in a refrigerated benchtop centrifuge at 4 °C. The clarified supernatant was divided into aliquots before being snap frozen in liquid nitrogen and stored at -80 °C.

Lysates were digested with 15 U of RNase I (Ambion; AM2294) per OD for 1 h at room temperature (25 °C), loaded onto 7.5-30% sucrose gradients, and spun at 41,000 rpm for 4 h and 45 min at 4 °C in an ultracentrifuge. These gradients offer better separation of the 40S peak than standard 10-50% gradients. The sucrose gradients were fractionated using a Brandel Density Gradient Fractionation System and the isolated 40S peaks were snap frozen in liquid nitrogen and stored at -80 °C.

RNA was purified from the 40S fractions using hot phenol chloroform, with an extended initial incubation of 1 h at 65 °C to reverse the formaldehyde cross-links. 15-80 nt ribosome footprints were size selected (see Table S8 for size standards) from a 15% TBE-Urea gel. Construction of 40S sequencing libraries was performed as described above for the RNA-seq sequencing libraries except that ribosomal RNA footprints were removed from the pooled ligated samples using the Ribo-Zero Gold rRNA Removal Kit Yeast (Illumina; MRZY1306) prior to reverse transcription.

### 80S ribosome profiling

#### Growth of yeast for 40S ribosome profiling

BY4741 (WT) and 6214 (*nam7*Δ [*upf1*Δ]) were grown in YPD. Cells were grown in 750 mL of media in a 2 L flask to a final OD_600_ of 0.6, fast filtered, and frozen in liquid nitrogen.

#### Preparation of 80S footprint libraries

Cells were lysed with a Retsch Cryomill (Retsch 20.749.0001) in the presence of frozen lysis buffer (20 mM Tris [pH 8], 140 mM KCl, 1.5 mM MgCl_2_, 1% Triton X-100) containing 0.1 mg/ml cycloheximide (Sigma; C7698). The milled cells were transferred to a 50 mL conical tube, thawed, and spun at 3000g for 5 min at 4 °C. The supernatant was transferred to 1.5 mL Eppendorf tubes and spun at full speed for 10 min in a refrigerated benchtop centrifuge at 4 °C. The clarified supernatant was divided into aliquots before being snap frozen in liquid nitrogen and stored at -80 °C. Construction of 80S sequencing libraries was performed as described above for the RNA-seq sequencing libraries except that Ribosomal RNA footprints were removed from the circularized libraries by oligonucleotide subtraction hybridization using Dynabeads™ MyOne™ Streptavidin C1 (Invitrogen; 65001) and a pool of DNA oligonucleotides that are the reverse complement of common rRNA contaminants (Table S8).

### Computational analysis

#### Read processing

Read analysis and sequence alignment were performed as previously described (Young *et al*., 2021). Fastq files (debarcoded by the core facility) were trimmed of their linkers and separated according to their 5-nt internal sample barcode by using CUTADAPT, and footprints with 7-nt UMIs measuring 57-77 nt, 32-41 nt, and 22-87 were retained for RNA-seq, 80S profiling, and 40S profiling samples respectively. Replicate datasets were combined to enhance sequencing depth. All PCR duplicates were removed using a simple python script to compare the 7-nt UMIs. The UMIs were removed with CUTADAPT and for RNA-seq samples only the first 50 nt of the trimmed read was retained. Contaminating tRNA and rRNA reads were removed with a BOWTIE alignment, allowing two mismatches, to a previously described improved index of noncoding RNAs (Young *et al*., 2021). Following removal of these ncRNA sequence, the resulting fastq files were aligned to the genome and then splice junctions using BOWTIE, allowing one mismatch but no multimapping reads. We used BOWTIE version 1.1.2 or 1.01 (Langmead et al., 2009) and included the parameter -y for all runs. We used the parameters -a -m 1 --best --strata for alignments to the genome and splice junctions.

Other analysis software used Biopython 1.58 or 1.63. In general, annotated ORFs marked dubious or those that overlapped with other transcripts were ignored in the analysis. Annotations for 5’UTRs and 3’ UTRs used were the most_abundant_transcripts in glucose media annotated in (Ng et al., 2020). We used coordinates from the R64-1-1 genome assembly from Saccharomyces Genome Database Project (Cherry et al., 2012). For detailed analysis of 5’UTRs, annotations from (Spealman et al., 2018) were used. Python code for the three basic analyses (Gene average, gene quantitation, and position average plots) has been published previously and are available here: www.github.com/guydoshlab/Yeastcode1

#### Analysis of alignments

Quantitation of gene-level footprint occupancy (i.e. counting reads mapping to genes for Figures S1b and c) was performed by creating footprint density in units of reads per kilobase per million mapped reads (rpkm) by taking the reads mapping to an annotated sequence (in rpm units) and dividing by the gene length in kilobases. To be included in the analysis, a threshold of 5 rpkm was required of ORF reads. 80S footprint reads were shifted by 13 nt and 40S footprint reads were shifted by 14 nt to correspond to the P site. mRNA-Seq reads were unshifted. Quantitation of peak reads around start or stop codons was performed by summing reads within a 5 nt region around the peak.

#### DESeq2 analysis

Listquant was used to calculate raw read counts for the coding sequence and introns of genes. Dubious ORFs were excluded from the analysis. Differential expression analysis was performed on reads from main open reading frames (ORFs) and introns with DESeq2 (Love *et al*., 2014). The two analyses were filtered using an FDR (padj) cutoff of 1% for ORFs and 5% for introns.

#### Pause score analysis

We calculated peak scores (also referred to as pause scores) for 5’UTR start codons from our *rpl11b*Δ *upf1*Δ 40S profiling data. The pause score is calculated as the number of reads at the motif (AUG) divided by the average number of reads within a window surrounding the motif (50nt to either side of the motif). 40S footprint reads were shifted by 13 nt to correspond to the P site. For comparison, we also computed peak heights in some cases, defined as the rpkm level of reads in the peak with a window.

For the 5’UTR AUG peak analysis start codons were identified in all frames from a custom genome annotation (GFF) file where the 5’UTR was extended either to the end of the next upstream ORF, or if the next upstream ORF was distant, a maximum length of 2kb. An alternative analysis utilizing a maximum length of 3 kb was used to detect outliers. One outlier, *FLO10*, extended beyond 3 kb and was hand annotated. Due to variability in 40S read coverage, which affected the denominator (average read window) of the pause score, raw peak height was used to detect 40S peaks. 40S peaks greater than 92 rpkm in height were regarded as true 40S peaks. Twenty-seven genes incorrectly assigned 5’UTR 40S peaks due to peaks on upstream unannotated transcripts and tRNAs were removed from the peak list. Three genes (*ASA1*, *LST7* and *PPT2*) had 0-frame peaks that lacked a stop codon between the AUG and the annotated start codon indicating that the detected 5’ peak was in fact a misannotated start codon (Table S9). These three genes with 0-frame peaks (*ASA1*, *LST7* and *PPT2*) were also removed for the list.

For the ORF peak analysis AUGs were identified in all frames within ORFs from the yeast genome annotation. 40S ORF peaks were filtered both by pause score (>10 fold) and peak height (>137 rpkm). This more stringent filtering was used for the ORF 40S peak analysis than for the 5’UTR 40S peak analysis due to the higher 40S background within ORFs caused by artifactual 80S ribosome separation during sample processing (Archer et al. 2016; Young et al. 2021). The 10-fold pause score cutoff was too stringent for three genes, *YOL159C-A*, *MAK31*, *KTI11*, due to the proximity of other 40S peaks resulting in an artificially elevated pause score denominator (average read window). The 40S peaks for these genes were added back to the 40S ORF peak list. (*YOL159C-A* - proximity of two out-of-frame 40S peaks, *MAK31* - proximity of an out-of-frame 40S peak with an internal start codon 40S peak, and *KTI11* - proximity of an out-of-frame 40S peak to both the canonical start codon 40S peak and an internal start codon 40S peak) Two additional peaks were added back for two genes, *INO4* and *PGA1* that had other peaks that passed the 10-fold pause score. (*INO4* - proximity of an out-of-frame 40S peak to a downstream out-of-frame 40S peak, and *PGA1* - proximity of the out-of-frame 40S peak to the canonical start codon) All twenty-two 0-frame 40S peaks were removed from the ORF 40S peak list that represented misannotated and internal start codons (Table S9). An ORF 40S peak representing a misannotated start codon was also detected for *LTO1*/*YNL260C*. This misannotation has been corrected in more recent yeast gffs and was removed from the ORF 40S peak list. Four of the genes that may also be misannotated start codons (*WSS1*, *YHL042W*, *AEP1*, and *PEX4*) had additional “ORF” 40S peaks upstream of the newly identified start codon (such as the old annotated start codon). These peaks were transferred to the 5’UTR 40S peak list as the newly identified start codons for these genes place these peaks within the newly established 5’UTR of these genes. One ORF AUG 40S peak for the gene *NQM1* was removed as it was caused by the proximity of an AUG to a stop codon (*AUG***UGA**).

### Analysis of related yeast sequences and sequencing data

For Figure 7C, DAL5 orthologs were identified by Blastn (v2.15.0+) (Camacho et al., 2009) using S. cerevisiae DAL5 as a query against seven genomes with parameters -word_size 7 -evalue 1e-2 -outfmt 6, and the top hit in each genome was considered the candidate ortholog. The protein sequences of DAL5 orthologs were then aligned using ClustalW in MEGA12 (Kumar et al., 2024) with default settings, and a phylogenetic tree was subsequently constructed using the neighbor-joining method (Saitou and Nei, 1987).

The mRNA-Seq from related yeast species (Blevins *et al*., 2019) was downloaded from Sequence Read Archive (SRA) with Project ID SRP187756. GFF annotations and genome references were found here: https://figshare.com/articles/dataset/Combination_of_novel_transcripts_from_de_novo_assembly_and_genes_from_reference_annotations_for_11_species/7851521/2

For *L. kluyveri*, the genome was obtained from the Saccharomyces Genome Database using the strain name NRRL Y-12651.

Raw bulk RNA-seq data were processed with fastp (v0.24.0) (Chen et al., 2018) to remove adapters, and the clean reads were aligned to the corresponding genomes using STAR (v2.7.11b) (Dobin et al., 2013) using default parameters except for outFilterMismatchNmax = 2 and alignEndsType = EndToEnd. BigWig (Kent et al., 2010) files were generated with bamCoverage from deepTools (v3.5.6) (Ramirez et al., 2014) using the parameters --binSize 10 --normalizeUsing BPM, and the tracks were visualized with IGV (Robinson et al., 2011). The codes for processing RNA-seq data can be accessed on Github: https://github.com/TriLab-bioinf/GUYDOSH_LAB_TK_207.

For Figure 7D (*S. pombe* data), RNA-Seq and ribosome profiling data from (Duncan and Mata, 2014) were processed the same. Reads were processed with: cutadapt -u 9 -a AAAAAAAA -m 15. Then, reads were aligned to a ncRNA to subtract reads that were not ribosomal footprints and then aligned to a spliced transcriptome as described in (Guydosh et al., 2017). Reads up to 50 nt were retained for mRNA-Seq and 25-34 nt for ribosome profiling. We used data files from ArrayExpress: E_pat1_Oh_mRNA and E_pat1_Oh_ribosome-protected RNA from E-MTAB-2179, which is starved for nitrogen. We used S_mRNA_wt_1A and S_ribo_wt_1A from E-MTAB-2176, which is grown in rich media.

### Northern blotting

Northern blotting was conducted using the DIG Northern Starter Kit (Roche; Cat. No. 12039672910).

#### Growth of cultures and RNA extraction for Northern blot

To determine if the 5’ uORFs identified by 40S ribosome profiling in the 5’-extended transcript of *DAL5* are responsible for triggering NMD, *dal5_ext*Δ *UPF1* and *dal5_ext*Δ *upf1*Δ cells transformed with the YCplac33-*DAL5* and YCplac33-*DAL5_noAUG* plasmids were grown in SC-U media to an OD_600_ of 1.0. The cells were harvested by being spun down at 4,000 rpm for 5 min at 4°C, the supernatant was discarded, and the cells frozen in liquid nitrogen. Similarly, for *DAL7*, *dal7_ext*Δ *UPF1* and *dal7_ext*Δ *upf1*Δ cells were transformed with the YCplac33-*DAL7* and YCplac33-*DAL7_noAUG* plasmids and were grown and harvested as described above for *DAL5*. To confirm that the *DAL5* and *DAL7* genes express two transcript isoforms, a 5’-extended transcript and a shorter induced canonical transcript (similar to LUTI 2-transcript systems), the above cells were also grown in poor nitrogen (proline) media to an OD_600_ of 1.0 and harvested as described above.

For the proline induction experiments, cells were grown in rich nitrogen (glutamine) media to an OD_600_ of 0.5 before the media was changed to poor nitrogen (proline). Samples were taken prior to induction, and at 0 m, 15 m, 30 m, 45 m, and 1 h after induction. Samples were processed as described above. For the glutamine repression experiments, cells were grown in poor nitrogen (proline) media to an OD_600_ of 0.5 before the media was changed to rich nitrogen (glutamine). Samples were taken prior to repression, and at 0 m, 5 m, 10 m, 20 m, and 1 h after induction. Samples were processed as described above.

Cells were resuspended in lysis buffer (8.4 mM EDTA, 60 mM Sodium acetate) and total RNA was purified using hot phenol chloroform.

#### Labeling of Northern probes

Single stranded antisense digoxigenin-labeled RNA probes were generated for *DAL5*, *DAL7*, and *SCR1* (loading control). To generate DNA templates for probe labeling, the last 500 nt of the *DAL5* and *DAL7* genes was PCR amplified using primer pairs DAL5_500nt_f/DAL5_probe_r and DAL7_500nt_f/DAL7_probe_r from the YCplac33-*DAL5* and YCplac33-*DAL7* plasmids. To generate a probe template for *SCR1* the last 500 nt of the *SCR1* gene was amplified by RT-PCR (SuperScript IV One-Step RT-PCR System with ezDNase; Invitrogen; Cat. No. 12595025) using primer pair SCR1_500nt_f and SCR1_probe_r from 1 μg of yeast total RNA. The reverse primers contain the T7 RNA Polymerase promoter sequence to allow transcription of antisense RNA probes. 200 ng of the purified probe templates were labeled in an in vitro transcription reaction with Digoxigenin-11-UTP according to the manufacturer’s instructions.

#### Formaldehyde gels

For *DAL5* northern blots 1 μg of denatured RNA/sample was loaded onto a 1.2 % agarose formaldehyde gel and run in 1x MOPS buffer at 80V for 2 h 30 min. For *DAL7* northern blots grown under poor nitrogen conditions a 2.5 % agarose was run for 3 h 30 m to ensure separation of the similarly sized 5’-extended and induced transcripts.

#### RNA transfer and fixation

RNA was transferred overnight to positively charged nylon membranes (Roche; Cat. No. 11209299001) using a Whatman Nytran SuPerCharge (SPC) TurboBlotter (Millipore Sigma Cat. No. WHA10416300). RNA was fixed to the membrane by UV crosslinking with a VWR UV crosslinker (VWR; Cat. No. 89131-484).

#### Hybridization and stringency washes

Membranes were hybridized in a VWR Hybridization Oven Model 5430 (VWR; Cat. No. 97005-252) at 68°C. Membranes were prehybridized in 10 mL of DIG Easy Hyb (Roche; 11796895001) for 30 min. The DIG-labeled probes (3.5 μL of 50 ng/μL) were denatured for 5 min at 100°C and then cooled on ice for 1 min before being added to 3.5 mL of DIG Easy Hyb. The membrane was hybridized with the probe/hybridization mixture for 6 h. The membrane was then washed twice in low stringency wash buffer (2x SSC, 0.1% SDS) at RT for 5 min. The membrane was then washed twice in prewarmed high stringency wash buffer (0.1x SSC, 0.1% SDS) at 68°C for 15 min.

#### Immunological detection

Immunological detection was carried out using buffers from the DIG Wash and Block Buffer Set (Roche; Cat. No. 11585762001). The membrane was washed in 1x Washing buffer for 5 min and then blocked in 1x Blocking solution for 30 min. The membrane was then incubated in antibody solution (Anti-Digoxigenin-AP [Roche; Cat. No. 11093274910] diluted 1:10,000 in 1x blocking buffer) for 30 min. The membrane was then washed twice in 1x Washing buffer for 15 min and equilibrated in Detection buffer for 5 min. The membrane was incubated with CDP-Star, ready-to-use solution (Roche; Cat. No. 12041677001) for 5 min before being imaged on an Amersham Imager 600.

#### Stripping and reprobing of RNA blots

Membranes were washed in RNase-free water for 5 min and then incubated in Stripping buffer (50 mM Tris-HCl, pH7.5, 5% SDS 50% Formamide) twice for 60 min at 80°C. The stripped membrane was washed twice in 2 x SSC at Rt for 5 min before reprobing with the *SCR1* loading control probe.

## REFERENCES

Albers, S., Allen, E.C., Bharti, N., Davyt, M., Joshi, D., Perez-Garcia, C.G., Santos, L., Mukthavaram, R., Delgado-Toscano, M.A., Molina, B., et al. (2023). Engineered tRNAs suppress nonsense mutations in cells and in vivo. Nature 618, 842–848. 10.1038/s41586-023-06133-1.

Amrani, N., Ganesan, R., Kervestin, S., Mangus, D.A., Ghosh, S., and Jacobson, A. (2004). A faux 3’-UTR promotes aberrant termination and triggers nonsense-mediated mRNA decay. Nature 432, 112–118. 10.1038/nature03060.

Andjus, S., Szachnowski, U., Vogt, N., Gioftsidi, S., Hatin, I., Cornu, D., Papadopoulos, C., Lopes, A., Namy, O., Wery, M., and Morillon, A. (2024). Pervasive translation of Xrn1-sensitive unstable long noncoding RNAs in yeast. RNA 30, 662–679. 10.1261/rna.079903.123.

Arribere, J.A., and Gilbert, W.V. (2013). Roles for transcript leaders in translation and mRNA decay revealed by transcript leader sequencing. Genome Res 23, 977–987. 10.1101/gr.150342.112.

Bharti, N., Santos, L., Davyt, M., Behrmann, S., Eichholtz, M., Jimenez-Sanchez, A., Hong, J.S., Rab, A., Sorscher, E.J., Albers, S., and Ignatova, Z. (2024). Translation velocity determines the efficacy of engineered suppressor tRNAs on pathogenic nonsense mutations. Nat Commun 15, 2957. 10.1038/s41467-024-47258-9.

Blevins, W.R., Carey, L.B., and Alba, M.M. (2019). Transcriptomics data of 11 species of yeast identically grown in rich media and oxidative stress conditions. BMC Res Notes 12, 250. 10.1186/s13104-019-4286-0.

Cai, H., Hauser, M., Naider, F., and Becker, J.M. (2007). Differential regulation and substrate preferences in two peptide transporters of Saccharomyces cerevisiae. Eukaryot Cell 6, 1805–1813. 10.1128/EC.00257-06.

Camacho, C., Coulouris, G., Avagyan, V., Ma, N., Papadopoulos, J., Bealer, K., and Madden, T.L. (2009). BLAST+: architecture and applications. BMC Bioinformatics 10, 421. 10.1186/1471-2105-10-421.

Celik, A., Baker, R., He, F., and Jacobson, A. (2017). High-resolution profiling of NMD targets in yeast reveals translational fidelity as a basis for substrate selection. RNA 23, 735–748. 10.1261/rna.060541.116.

Chen, J., Tresenrider, A., Chia, M., McSwiggen, D.T., Spedale, G., Jorgensen, V., Liao, H., van Werven, F.J., and Unal, E. (2017). Kinetochore inactivation by expression of a repressive mRNA. Elife 6. 10.7554/eLife.27417.

Chen, S., Zhou, Y., Chen, Y., and Gu, J. (2018). fastp: an ultra-fast all-in-one FASTQ preprocessor. Bioinformatics 34, i884–i890. 10.1093/bioinformatics/bty560.

Cheng, Z., Otto, G.M., Powers, E.N., Keskin, A., Mertins, P., Carr, S.A., Jovanovic, M., and Brar, G.A. (2018). Pervasive, Coordinated Protein-Level Changes Driven by Transcript Isoform Switching during Meiosis. Cell 172, 910–923 e916. 10.1016/j.cell.2018.01.035.

Cherry, J.M., Hong, E.L., Amundsen, C., Balakrishnan, R., Binkley, G., Chan, E.T., Christie, K.R., Costanzo, M.C., Dwight, S.S., Engel, S.R., et al. (2012). Saccharomyces Genome Database: the genomics resource of budding yeast. Nucleic Acids Res 40, D700–705. 10.1093/nar/gkr1029.

Chia, M., Tresenrider, A., Chen, J., Spedale, G., Jorgensen, V., Unal, E., and van Werven, F.J. (2017). Transcription of a 5’ extended mRNA isoform directs dynamic chromatin changes and interference of a downstream promoter. Elife 6. 10.7554/eLife.27420.

Coelho, J.P.L., Yip, M.C.J., Oltion, K., Taunton, J., and Shao, S. (2024). The eRF1 degrader SRI-41315 acts as a molecular glue at the ribosomal decoding center. Nat Chem Biol 20, 877–884. 10.1038/s41589-023-01521-0.

Coller, J., and Ignatova, Z. (2024). tRNA therapeutics for genetic diseases. Nat Rev Drug Discov 23, 108–125. 10.1038/s41573-023-00829-9.

Cui, Y., Hagan, K.W., Zhang, S., and Peltz, S.W. (1995). Identification and characterization of genes that are required for the accelerated degradation of mRNAs containing a premature translational termination codon. Genes Dev 9, 423–436. 10.1101/gad.9.4.423.

Decourty, L., Doyen, A., Malabat, C., Frachon, E., Rispal, D., Seraphin, B., Feuerbach, F., Jacquier, A., and Saveanu, C. (2014). Long open reading frame transcripts escape nonsense-mediated mRNA decay in yeast. Cell Rep 6, 593–598. 10.1016/j.celrep.2014.01.025.

Dehecq, M., Decourty, L., Namane, A., Proux, C., Kanaan, J., Le Hir, H., Jacquier, A., and Saveanu, C. (2018). Nonsense-mediated mRNA decay involves two distinct Upf1-bound complexes. EMBO J 37. 10.15252/embj.201899278.

Dobin, A., Davis, C.A., Schlesinger, F., Drenkow, J., Zaleski, C., Jha, S., Batut, P., Chaisson, M., and Gingeras, T.R. (2013). STAR: ultrafast universal RNA-seq aligner. Bioinformatics 29, 15–21. 10.1093/bioinformatics/bts635.

Duncan, C.D., and Mata, J. (2014). The translational landscape of fission-yeast meiosis and sporulation. Nat Struct Mol Biol 21, 641–647. 10.1038/nsmb.2843.

Farabaugh, P.J., Kramer, E., Vallabhaneni, H., and Raman, A. (2006). Evolution of +1 programmed frameshifting signals and frameshift-regulating tRNAs in the order Saccharomycetales. J Mol Evol 63, 545–561. 10.1007/s00239-005-0311-0.

Fritz, S.E., Ranganathan, S., Wang, C.D., and Hogg, J.R. (2020). The RNA-binding protein PTBP1 promotes ATPase-dependent dissociation of the RNA helicase UPF1 to protect transcripts from nonsense-mediated mRNA decay. J Biol Chem 295, 11613–11625. 10.1074/jbc.RA120.013824.

Gaba, A., Jacobson, A., and Sachs, M.S. (2005). Ribosome occupancy of the yeast CPA1 upstream open reading frame termination codon modulates nonsense-mediated mRNA decay. Mol Cell 20, 449–460. 10.1016/j.molcel.2005.09.019.

Geisler, S., Lojek, L., Khalil, A.M., Baker, K.E., and Coller, J. (2012). Decapping of long noncoding RNAs regulates inducible genes. Mol Cell 45, 279–291. 10.1016/j.molcel.2011.11.025.

Goetz, A.E., and Wilkinson, M. (2017). Stress and the nonsense-mediated RNA decay pathway. Cell Mol Life Sci 74, 3509–3531. 10.1007/s00018-017-2537-6.

Gurzeler, L.A., Link, M., Ibig, Y., Schmidt, I., Galuba, O., Schoenbett, J., Gasser-Didierlaurant, C., Parker, C.N., Mao, X., Bitsch, F., et al. (2023). Drug-induced eRF1 degradation promotes readthrough and reveals a new branch of ribosome quality control. Cell Rep 42, 113056. 10.1016/j.celrep.2023.113056.

Guydosh, N.R., Kimmig, P., Walter, P., and Green, R. (2017). Regulated Ire1-dependent mRNA decay requires no-go mRNA degradation to maintain endoplasmic reticulum homeostasis in S. pombe. Elife 6. 10.7554/eLife.29216.

He, F., and Jacobson, A. (1995). Identification of a novel component of the nonsense-mediated mRNA decay pathway by use of an interacting protein screen. Genes Dev 9, 437–454. 10.1101/gad.9.4.437.

He, F., and Jacobson, A. (2015). Nonsense-Mediated mRNA Decay: Degradation of Defective Transcripts Is Only Part of the Story. Annu Rev Genet 49, 339–366. 10.1146/annurev-genet-112414-054639.

He, F., Li, X., Spatrick, P., Casillo, R., Dong, S., and Jacobson, A. (2003). Genome-wide analysis of mRNAs regulated by the nonsense-mediated and 5’ to 3’ mRNA decay pathways in yeast. Mol Cell 12, 1439–1452. 10.1016/s1097-2765(03)00446-5.

He, F., Peltz, S.W., Donahue, J.L., Rosbash, M., and Jacobson, A. (1993). Stabilization and ribosome association of unspliced pre-mRNAs in a yeast upf1-mutant. Proc Natl Acad Sci U S A 90, 7034–7038. 10.1073/pnas.90.15.7034.

Hogg, J.R., and Goff, S.P. (2010). Upf1 senses 3’UTR length to potentiate mRNA decay. Cell 143, 379–389. 10.1016/j.cell.2010.10.005.

Houseley, J., Rubbi, L., Grunstein, M., Tollervey, D., and Vogelauer, M. (2008). A ncRNA modulates histone modification and mRNA induction in the yeast GAL gene cluster. Mol Cell 32, 685–695. 10.1016/j.molcel.2008.09.027.

Johansson, M.J., He, F., Spatrick, P., Li, C., and Jacobson, A. (2007). Association of yeast Upf1p with direct substrates of the NMD pathway. Proc Natl Acad Sci U S A 104, 20872–20877. 10.1073/pnas.0709257105.

Kebaara, B.W., and Atkin, A.L. (2009). Long 3’-UTRs target wild-type mRNAs for nonsense-mediated mRNA decay in Saccharomyces cerevisiae. Nucleic Acids Res 37, 2771–2778. 10.1093/nar/gkp146.

Kent, W.J., Zweig, A.S., Barber, G., Hinrichs, A.S., and Karolchik, D. (2010). BigWig and BigBed: enabling browsing of large distributed datasets. Bioinformatics 26, 2204–2207. 10.1093/bioinformatics/btq351.

Kim, J.H., Lee, B.B., Oh, Y.M., Zhu, C., Steinmetz, L.M., Lee, Y., Kim, W.K., Lee, S.B., Buratowski, S., and Kim, T. (2016). Modulation of mRNA and lncRNA expression dynamics by the Set2-Rpd3S pathway. Nat Commun 7, 13534. 10.1038/ncomms13534.

Kim, T., Xu, Z., Clauder-Munster, S., Steinmetz, L.M., and Buratowski, S. (2012). Set3 HDAC mediates effects of overlapping noncoding transcription on gene induction kinetics. Cell 150, 1158–1169. 10.1016/j.cell.2012.08.016.

Kishor, A., Ge, Z., and Hogg, J.R. (2019). hnRNP L-dependent protection of normal mRNAs from NMD subverts quality control in B cell lymphoma. EMBO J 38. 10.15252/embj.201899128.

Kolakada, D., Fu, R., Biziaev, N., Shuvalov, A., Lore, M., Campbell, A.E., Cortazar, M.A., Sajek, M.P., Hesselberth, J.R., Mukherjee, N., et al. (2025). Systematic analysis of nonsense variants uncovers peptide release rate as a novel modifier of nonsense-mediated mRNA decay. Cell Genom 5, 100882. 10.1016/j.xgen.2025.100882.

Kumar, S., Stecher, G., Suleski, M., Sanderford, M., Sharma, S., and Tamura, K. (2024). MEGA12: Molecular Evolutionary Genetic Analysis Version 12 for Adaptive and Green Computing. Mol Biol Evol 41. 10.1093/molbev/msae263.

Langmead, B., Trapnell, C., Pop, M., and Salzberg, S.L. (2009). Ultrafast and memory-efficient alignment of short DNA sequences to the human genome. Genome Biol 10, R25. 10.1186/gb-2009-10-3-r25.

Le Hir, H., Gatfield, D., Izaurralde, E., and Moore, M.J. (2001). The exon-exon junction complex provides a binding platform for factors involved in mRNA export and nonsense-mediated mRNA decay. EMBO J 20, 4987–4997. 10.1093/emboj/20.17.4987.

Lee, B.S., and Culbertson, M.R. (1995). Identification of an additional gene required for eukaryotic nonsense mRNA turnover. Proc Natl Acad Sci U S A 92, 10354–10358. 10.1073/pnas.92.22.10354.

Leeds, P., Peltz, S.W., Jacobson, A., and Culbertson, M.R. (1991). The product of the yeast UPF1 gene is required for rapid turnover of mRNAs containing a premature translational termination codon. Genes Dev 5, 2303–2314. 10.1101/gad.5.12a.2303.

Leeds, P., Wood, J.M., Lee, B.S., and Culbertson, M.R. (1992). Gene products that promote mRNA turnover in Saccharomyces cerevisiae. Mol Cell Biol 12, 2165–2177. 10.1128/mcb.12.5.2165-2177.1992.

Losson, R., and Lacroute, F. (1979). Interference of nonsense mutations with eukaryotic messenger RNA stability. Proc Natl Acad Sci U S A 76, 5134–5137. 10.1073/pnas.76.10.5134.

Love, M.I., Huber, W., and Anders, S. (2014). Moderated estimation of fold change and dispersion for RNA-seq data with DESeq2. Genome Biol 15, 550. 10.1186/s13059-014-0550-8.

Luke, B., Azzalin, C.M., Hug, N., Deplazes, A., Peter, M., and Lingner, J. (2007). Saccharomyces cerevisiae Ebs1p is a putative ortholog of human Smg7 and promotes nonsense-mediated mRNA decay. Nucleic Acids Res 35, 7688–7697. 10.1093/nar/gkm912.

Mabin, J.W., Vock, I.W., Machyna, M., Haque, N., Thakran, P., Zhang, A., Rai, G., Leibler, I.N., Inglese, J., Simon, M.D., and Hogg, J.R. (2025). Uncovering the isoform-resolution kinetic landscape of nonsense-mediated mRNA decay with EZbakR. bioRxiv. 10.1101/2025.03.12.642874.

Malabat, C., Feuerbach, F., Ma, L., Saveanu, C., and Jacquier, A. (2015). Quality control of transcription start site selection by nonsense-mediated-mRNA decay. Elife 4. 10.7554/eLife.06722.

Mangkalaphiban, K., He, F., Ganesan, R., Wu, C., Baker, R., and Jacobson, A. (2021). Transcriptome-wide investigation of stop codon readthrough in Saccharomyces cerevisiae. PLoS Genet 17, e1009538. 10.1371/journal.pgen.1009538.

Maquat, L.E., Kinniburgh, A.J., Rachmilewitz, E.A., and Ross, J. (1981). Unstable beta-globin mRNA in mRNA-deficient beta o thalassemia. Cell 27, 543–553. 10.1016/0092-8674(81)90396-2.

Martens, J.A., Laprade, L., and Winston, F. (2004). Intergenic transcription is required to repress the Saccharomyces cerevisiae SER3 gene. Nature 429, 571–574. 10.1038/nature02538.

May, G.E., Akirtava, C., Agar-Johnson, M., Micic, J., Woolford, J., and McManus, J. (2023). Unraveling the influences of sequence and position on yeast uORF activity using massively parallel reporter systems and machine learning. Elife 12. 10.7554/eLife.69611.

McGlincy, N.J., and Ingolia, N.T. (2017). Transcriptome-wide measurement of translation by ribosome profiling. Methods 126, 112–129. 10.1016/j.ymeth.2017.05.028.

Meydan, S., and Guydosh, N.R. (2020). Disome and Trisome Profiling Reveal Genome-wide Targets of Ribosome Quality Control. Mol Cell 79, 588–602 e586. 10.1016/j.molcel.2020.06.010.

Miura, F., Kawaguchi, N., Sese, J., Toyoda, A., Hattori, M., Morishita, S., and Ito, T. (2006). A large-scale full-length cDNA analysis to explore the budding yeast transcriptome. Proc Natl Acad Sci U S A 103, 17846–17851. 10.1073/pnas.0605645103.

Morse, K., Bishop, A.L., Swerdlow, S., Leslie, J.M., and Unal, E. (2024). Swi/Snf chromatin remodeling regulates transcriptional interference and gene repression. Mol Cell 84, 3080–3097 e3089. 10.1016/j.molcel.2024.06.029.

Mort, M., Ivanov, D., Cooper, D.N., and Chuzhanova, N.A. (2008). A meta-analysis of nonsense mutations causing human genetic disease. Hum Mutat 29, 1037–1047. 10.1002/humu.20763.

Muhlrad, D., and Parker, R. (1999). Aberrant mRNAs with extended 3’ UTRs are substrates for rapid degradation by mRNA surveillance. RNA 5, 1299–1307. 10.1017/s1355838299990829.

Munoz, O., Lore, M., and Jagannathan, S. (2023). The long and short of EJC-independent nonsense-mediated RNA decay. Biochem Soc Trans 51, 1121–1129. 10.1042/BST20221131.

Nagalakshmi, U., Wang, Z., Waern, K., Shou, C., Raha, D., Gerstein, M., and Snyder, M. (2008). The transcriptional landscape of the yeast genome defined by RNA sequencing. Science 320, 1344–1349. 10.1126/science.1158441.

Ng, P.C., Wong, E.D., MacPherson, K.A., Aleksander, S., Argasinska, J., Dunn, B., Nash, R.S., Skrzypek, M.S., Gondwe, F., Jha, S., et al. (2020). Transcriptome visualization and data availability at the Saccharomyces Genome Database. Nucleic Acids Res 48, D743–D748. 10.1093/nar/gkz892.

Oliveira, C.C., and McCarthy, J.E. (1995). The relationship between eukaryotic translation and mRNA stability. A short upstream open reading frame strongly inhibits translational initiation and greatly accelerates mRNA degradation in the yeast Saccharomyces cerevisiae. J Biol Chem 270, 8936–8943. 10.1074/jbc.270.15.8936.

Oltion, K., Carelli, J.D., Yang, T., See, S.K., Wang, H.Y., Kampmann, M., and Taunton, J. (2023). An E3 ligase network engages GCN1 to promote the degradation of translation factors on stalled ribosomes. Cell 186, 346–362 e317. 10.1016/j.cell.2022.12.025.

Pacheco, M., D’Orazio, K.N., Lessen, L.N., Veltri, A.J., Neiman, Z., Loll-Krippleber, R., Brown, G.W., and Green, R. (2024). Genetic screens in Saccharomyces cerevisiae identify a role for 40S ribosome recycling factors Tma20 and Tma22 in nonsense-mediated decay. G3 (Bethesda) 14. 10.1093/g3journal/jkad295.

Palanimurugan, R., Scheel, H., Hofmann, K., and Dohmen, R.J. (2004). Polyamines regulate their synthesis by inducing expression and blocking degradation of ODC antizyme. EMBO J 23, 4857–4867. 10.1038/sj.emboj.7600473.

Peccarelli, M., Scott, T.D., Steele, M., and Kebaara, B.W. (2016). mRNAs involved in copper homeostasis are regulated by the nonsense-mediated mRNA decay pathway depending on environmental conditions. Fungal Genet Biol 86, 81–90. 10.1016/j.fgb.2015.12.011.

Pelechano, V., Wei, W., and Steinmetz, L.M. (2013). Extensive transcriptional heterogeneity revealed by isoform profiling. Nature 497, 127–131. 10.1038/nature12121.

Peltz, S.W., Brown, A.H., and Jacobson, A. (1993). mRNA destabilization triggered by premature translational termination depends on at least three cis-acting sequence elements and one trans-acting factor. Genes Dev 7, 1737–1754. 10.1101/gad.7.9.1737.

Peltz, S.W., Morsy, M., Welch, E.M., and Jacobson, A. (2013). Ataluren as an agent for therapeutic nonsense suppression. Annu Rev Med 64, 407–425. 10.1146/annurev-med-120611-144851.

Poonia, P., Valabhoju, V., Li, T., Iben, J., Niu, X., Lin, Z., and Hinnebusch, A.G. (2025). Yeast poly(A)-binding protein (Pab1) controls translation initiation in vivo primarily by blocking mRNA decapping and decay. Nucleic Acids Res 53. 10.1093/nar/gkaf143.

Porrua, O., Hobor, F., Boulay, J., Kubicek, K., D’Aubenton-Carafa, Y., Gudipati, R.K., Stefl, R., and Libri, D. (2012). In vivo SELEX reveals novel sequence and structural determinants of Nrd1-Nab3-Sen1-dependent transcription termination. EMBO J 31, 3935–3948. 10.1038/emboj.2012.237.

Rai, R., Genbauffe, F.S., and Cooper, T.G. (1988). Structure and transcription of the allantoate permease gene (DAL5) from Saccharomyces cerevisiae. J Bacteriol 170, 266–271. 10.1128/jb.170.1.266-271.1988.

Ramirez, F., Dundar, F., Diehl, S., Gruning, B.A., and Manke, T. (2014). deepTools: a flexible platform for exploring deep-sequencing data. Nucleic Acids Res 42, W187–191. 10.1093/nar/gku365.

Robinson, J.T., Thorvaldsdottir, H., Winckler, W., Guttman, M., Lander, E.S., Getz, G., and Mesirov, J.P. (2011). Integrative genomics viewer. Nat Biotechnol 29, 24–26. 10.1038/nbt.1754.

Ruiz-Echevarria, M.J., and Peltz, S.W. (2000). The RNA binding protein Pub1 modulates the stability of transcripts containing upstream open reading frames. Cell 101, 741–751. 10.1016/s0092-8674(00)80886-7.

Rutherford, K.M., Lera-Ramirez, M., and Wood, V. (2024). PomBase: a Global Core Biodata Resource-growth, collaboration, and sustainability. Genetics 227. 10.1093/genetics/iyae007.

Saitou, N., and Nei, M. (1987). The neighbor-joining method: a new method for reconstructing phylogenetic trees. Mol Biol Evol 4, 406–425. 10.1093/oxfordjournals.molbev.a040454.

Sharma, J., Du, M., Wong, E., Mutyam, V., Li, Y., Chen, J., Wangen, J., Thrasher, K., Fu, L., Peng, N., et al. (2021). A small molecule that induces translational readthrough of CFTR nonsense mutations by eRF1 depletion. Nat Commun 12, 4358. 10.1038/s41467-021-24575-x.

Singh, G., Rebbapragada, I., and Lykke-Andersen, J. (2008). A competition between stimulators and antagonists of Upf complex recruitment governs human nonsense-mediated mRNA decay. PLoS Biol 6, e111. 10.1371/journal.pbio.0060111.

Smith, J.E., Alvarez-Dominguez, J.R., Kline, N., Huynh, N.J., Geisler, S., Hu, W., Coller, J., and Baker, K.E. (2014). Translation of small open reading frames within unannotated RNA transcripts in Saccharomyces cerevisiae. Cell Rep 7, 1858–1866. 10.1016/j.celrep.2014.05.023.

Spealman, P., Naik, A.W., May, G.E., Kuersten, S., Freeberg, L., Murphy, R.F., and McManus, J. (2018). Conserved non-AUG uORFs revealed by a novel regression analysis of ribosome profiling data. Genome Res 28, 214–222. 10.1101/gr.221507.117.

ter Schure, E.G., van Riel, N.A., and Verrips, C.T. (2000). The role of ammonia metabolism in nitrogen catabolite repression in Saccharomyces cerevisiae. FEMS Microbiol Rev 24, 67–83. 10.1111/j.1574-6976.2000.tb00533.x.

Tresenrider, A., Morse, K., Jorgensen, V., Chia, M., Liao, H., van Werven, F.J., and Unal, E. (2021). Integrated genomic analysis reveals key features of long undecoded transcript isoform-based gene repression. Mol Cell 81, 2231–2245 e2211. 10.1016/j.molcel.2021.03.013.

Van Dalfsen, K.M., Hodapp, S., Keskin, A., Otto, G.M., Berdan, C.A., Higdon, A., Cheunkarndee, T., Nomura, D.K., Jovanovic, M., and Brar, G.A. (2018). Global Proteome Remodeling during ER Stress Involves Hac1-Driven Expression of Long Undecoded Transcript Isoforms. Dev Cell 46, 219–235 e218. 10.1016/j.devcel.2018.06.016.

van Dijk, E.L., Chen, C.L., d’Aubenton-Carafa, Y., Gourvennec, S., Kwapisz, M., Roche, V., Bertrand, C., Silvain, M., Legoix-Ne, P., Loeillet, S., et al. (2011). XUTs are a class of Xrn1-sensitive antisense regulatory non-coding RNA in yeast. Nature 475, 114–117. 10.1038/nature10118.

Wery, M., Descrimes, M., Vogt, N., Dallongeville, A.S., Gautheret, D., and Morillon, A. (2016). Nonsense-Mediated Decay Restricts LncRNA Levels in Yeast Unless Blocked by Double-Stranded RNA Structure. Mol Cell 61, 379–392. 10.1016/j.molcel.2015.12.020.

Wu, C., Roy, B., He, F., Yan, K., and Jacobson, A. (2020). Poly(A)-Binding Protein Regulates the Efficiency of Translation Termination. Cell Rep 33, 108399. 10.1016/j.celrep.2020.108399.

Xu, Z., Wei, W., Gagneur, J., Perocchi, F., Clauder-Munster, S., Camblong, J., Guffanti, E., Stutz, F., Huber, W., and Steinmetz, L.M. (2009). Bidirectional promoters generate pervasive transcription in yeast. Nature 457, 1033–1037. 10.1038/nature07728.

Yepiskoposyan, H., Aeschimann, F., Nilsson, D., Okoniewski, M., and Muhlemann, O. (2011). Autoregulation of the nonsense-mediated mRNA decay pathway in human cells. RNA 17, 2108–2118. 10.1261/rna.030247.111.

Young, D.J., Meydan, S., and Guydosh, N.R. (2021). 40S ribosome profiling reveals distinct roles for Tma20/Tma22 (MCT-1/DENR) and Tma64 (eIF2D) in 40S subunit recycling. Nat Commun 12, 2976. 10.1038/s41467-021-23223-8.

Zhang, Z., and Dietrich, F.S. (2005). Mapping of transcription start sites in Saccharomyces cerevisiae using 5’ SAGE. Nucleic Acids Res 33, 2838–2851. 10.1093/nar/gki583.

